# PDLIM5 Modulates YAP1 Localisation and Fibrogenic Gene Expression in Hepatic Stellate Cells

**DOI:** 10.64898/2026.09.25.748507

**Authors:** Kara Worsley, Dina Abdelmottaleb, Lana Ibrahim, Elliot F Jennings, Hayato Kakinuma, Joel Malungu Makopa, Anna Meier, Rabea Pätzold, Lindsay Birchall, Varinder S Athwal, Elliot Jokl, Tristan Mckay, Karen Piper Hanley, Jonathan D. Humphries, James Pritchett

## Abstract

Hepatic stellate cells (HSCs) are the key cellular drivers of liver fibrosis. During liver injury and chronic inflammation HSCs adopt an activated phenotype and secrete fibrotic extracellular matrix (ECM) components such as collagen I. Mechanical cues derived from the fibrotic ECM drive and support the activation of HSCs, via mechanisms that involve integrins and the mechano-sensitive transcriptional regulator YAP1. It is not yet well understood how external mechanical cues are translated into a molecular response that alters YAP1 nuclear shuttling. There is evidence that suggests the PDZ and LIM domain protein (PDLIM) 5 can regulate YAP1 shuttling in human epithelial cells. We therefore investigated whether PDLIM5 is expressed in HSCs and contributes to YAP1 associated HSC mechano-activation in the context of liver fibrosis. PDLIM5 protein was detected in HSCs in fibrotic human and mouse liver. PDLIM5 transcript and protein were expressed by primary human and mouse HSCs and by the immortalised HSC LX-2 cell line. PDLIM5 colocalised with integrin adhesion complexes at actin stress fibres suggesting a role in HSC ECM adhesion. Co-immunoprecipitation and proximity ligation in LX-2 cells support an association between PDLIM5 and YAP1. We used pharmacological (paclitaxel) and genetic (siRNA and CRISPRi) approaches to inhibit PDLIM5 in HSCs. Inhibiting PDLIM5 reduced YAP1 nuclear localisation and fibrotic gene (*COL1A1*, *ACTA2*) expression in LX-2 cells. Overall, these data support a role for PDLIM5 in regulating YAP1 localisation and fibrogenic gene expression in HSCs and suggest that PDLIM5 may influence the development and maintenance of liver fibrosis.

## INTRODUCTION

Cells interact with their external environment via membrane spanning integrins^1^. Integrins are a family of extracellular matrix (ECM) receptors that associate with dynamic signalling complexes called focal adhesions, and have been investigated as drug targets in liver fibrosis^2^. Integrins, focal adhesion proteins (e.g. paxillin) and associated proteins form the adhesome^3,4^. Mechanical cues such as shear (e.g. blood flow) or changes in substrate stiffness influence cell behaviour via mechano-signalling pathways modulated by integrins and the other components of the adhesome. These mechano-signalling processes have been implicated as key drivers of organ fibrosis^5,6^, including in the liver^7–10^.

Injury to the liver causes an inflammatory response that drives the activation of hepatic stellate cells (HSCs)^11^. HSCs are well established as the cellular drivers of liver fibrosis^12–14^. HSCs adopt an activated myofibroblast phenotype in response to liver injury. The activated HSC phenotype is associated with increased production of fibrotic extracellular matrix (ECM) components such as collagen I^12^. The fibrotic environment has increased mechanical stiffness^15^ which can be measured for non-invasive diagnosis and staging of liver fibrosis^16^. Importantly, increased mechanical stiffness is necessary for the initial activation of HSCs and for maintenance of the activated myofibroblast phenotype^17–20^. Yes-associated protein 1 (YAP1) is a mechano-sensitive transcriptional regulator^21^ which promotes HSC activation in response to elevated substrate stiffness^8,9^. YAP1 activates target gene expression via interaction with the TEAD family of transcription factors^22–24^. Inhibition of the interaction between YAP1 and TEAD using verteporfin^25^ reduced HSC activation *in vitro* and liver fibrosis *in vivo*^8^. It remains unclear how external mechanical cues translated via the cytoskeleton regulate YAP1 activity. In an epithelial cell line PDZ and LIM Domain (PDLIM) enigma family proteins PDLIM5 and 7 have been implicated in the mechano- regulation of YAP1 activity^26^. This mechanism could provide an explanation for YAP1 mediated mechano-activation of HSCs during liver fibrosis. Intriguingly two of our previous datasets comparing global gene expression in quiescent and *in vitro* activated primary HSCs showed *PDLIM5* was significantly upregulated during HSC activation^8,27^. Furthermore, ECM interactions with integrins, including α11β1^8^, regulate HSC phenotype and PDLIM5 and 7 are components of the integrin adhesome^3,4^. Despite this, the function of PDLIM family proteins during HSC mechano-activation remains to be elucidated. Here we demonstrate that PDLIM5 is expressed by HSCs, and data further suggest that PDLIM5 interacts with YAP1 and regulates YAP1 activity in the context of HSC mechano-activation.

## RESULTS

### PDLIM5 is expressed by HSCs during liver fibrosis *in vivo*

Our previous gene expression datasets showed increased *PDLIM5* expression during *in vitro* activation of mouse primary HSCs^8,27^. Based on this observation we investigated the potential for PDLIM5 to be involved in YAP1 mediated mechano-activation of HSCs. In support of a potential role in fibrogenesis, IHC was performed and PDLIM5 was detected in both human and experimentally induced mouse liver fibrosis. In fibrotic liver, PDLIM5 expression was observed in spindle-shaped cells associated with fibrotic regions, morphologically consistent with activated hepatic stellate cells (**Figure 1A and B**). Sirius red staining of an equivalent portal region from a near serial section is shown in supplemental figure 1. In normal mouse liver, PDLIM5 staining was readily observed in the cytoplasm and near the membrane of hepatocytes (**Figure 1B**). The localisation of PDLIM5 in hepatocytes appeared to be more membrane associated in both fibrotic human (**Figure 1A**) and experimental mouse fibrosis (**Figure 1B**). Haematoxylin and eosin, and Sirius red of an equivalent region in near serial sections of experimental mouse fibrosis are shown in supplemental figure 2. Dual immunofluorescence confirmed expression of PDLIM5 in ACTA2-positive cells within human fibrotic scar tissue (**Figure 2**). Furthermore, data from two independent singe cell RNA sequencing datasets^28,29^ indicates that there is a population of *ACTA2* positive cells in the human liver that express *PDLIM5* (**supplemental figure 3 and 4**). We suggest these are likely to be *PDLIM5* positive HSCs. Together this data supports a potential role for PDLIM5 during HSC activation

**Figure 1.**
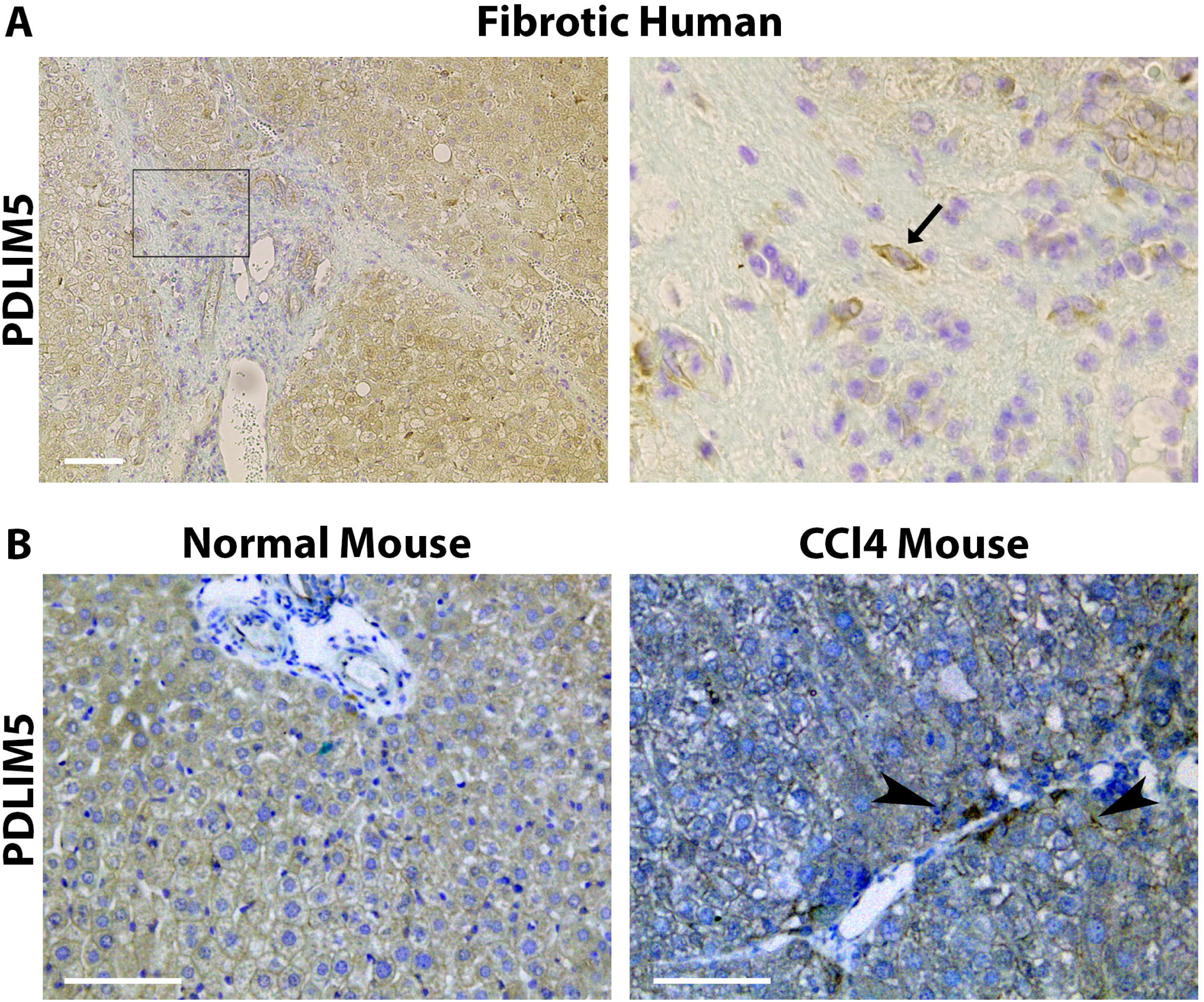
PDLIM5 (brown) counterstained with toludine blue. **A**. Fibrotic human liver. Region in box (left panel) is shown at higher magnification in right panel. Arrow shows example of HSC positive for PDLIM5. **B**. Normal (olive oil control, left) and CCl4-induced fibrosis (right) mouse liver. Arrow heads in CCl4 mouse indicate PDLIM5 positive HSCs. Scale bars are 100µm.

**Figure 2.**
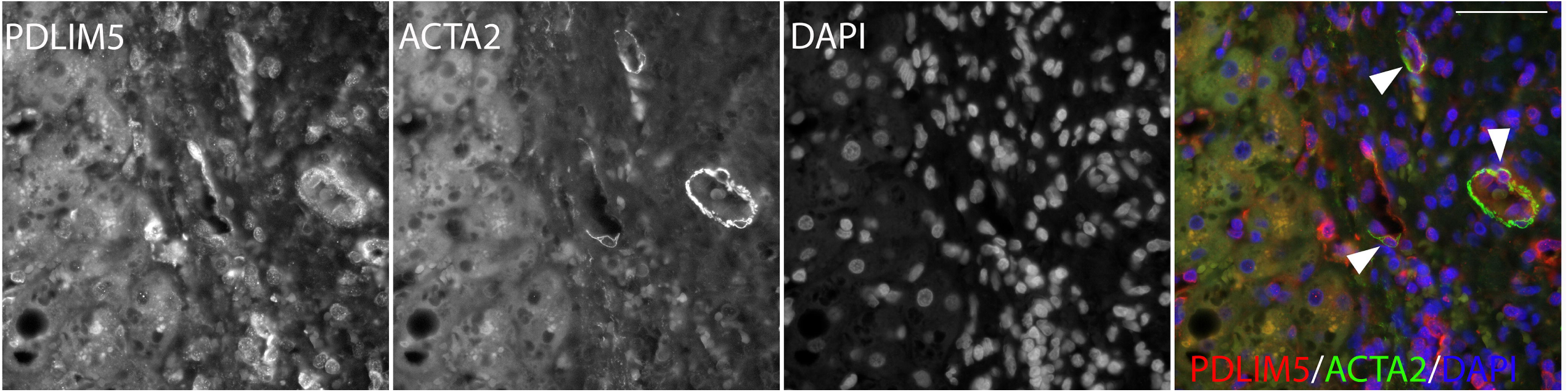
Human fibrotic liver. PDLIM5 (red) is present in ACTA2 (green) positive cells (arrowheads). Scale bar is 50µm.

### PDLIM5 is Expressed by HSCs *in vitro*

PDLIM5 transcript and protein were detected in both immortalised (**Figure 3A**) and primary HSCs (**Figure 3B**). *PDLIM5* was significantly (*p<0.05*) upregulated during culture activation of mHSCs (**Figure 3B**). In contrast in LX-2 cells treatment with TGFb did not change *PDLIM5* expression despite promotion of fibrotic gene expression (**Figure 3A**). Protein expression of PDLIM5 in HSCs was confirmed by western blot of LX-2 and mHSC lysates (**Figure 3C and 3D**), but in LX-2 cells TGFb did not appear to alter PDLIM5 protein expression, despite increased expression during culture activation of primary HSCs. These observations suggest that PDLIM5 expression may be more closely associated with HSC activation and/or mechanical context than regulation by TGFb signalling.

**Figure 3.**
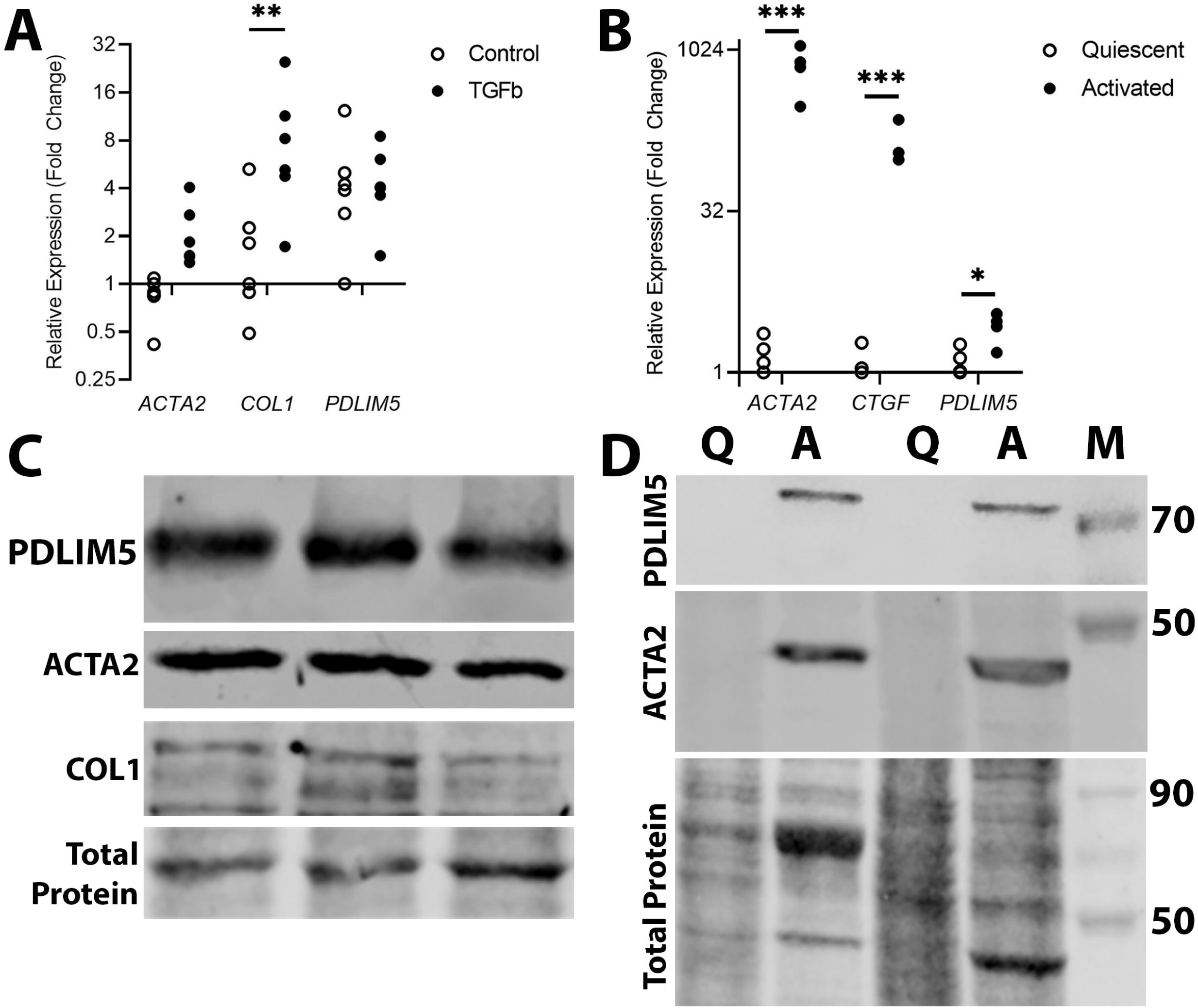
HSCs Express PDLIM5. **A**. qPCR of control and TGFb treated LX-2 cells, reference gene *GusB* (*n=6, *p<0.05).* **B**. qPCR comparing quiescent and culture activated mHSCs (*n=3*, *** p<0.001, *\* p<0.05)*. **C**. LX-2 cells express PDLIM5 protein, ACTA2, and COL1. **D**. Culture activated primary mHSCs express PDLIM5 protein and ACTA2. Q, quiescent; A, activated. M, Marker.

### Subcellular PDLIM5 Protein Localisation in HSCs

PDLIM5 was expressed by ACTA2 positive mHSCs **(Figure 4A**). PDLIM5 protein was not uniformly distributed throughout the mHSC cytoplasm but instead appeared to be enriched at ACTA2 positive structures at the cell periphery **(Figure 4A**). YAP1 was partially co-localised with PDLIM5 protein in the cytoplasm of mouse (**Figure 4B**) and human (**Figure 4C**) primary HSCs. In primary mouse and human HSCs (**Figure 4B and C**) YAP1 was predominantly nuclear, as reported previously in mHSCs^8,9^. The pattern of YAP1 and PDLIM5 co-localisation supports a potential role for PDLIM5 linking adhesion-associated structures and YAP1 signalling.

**Figure 4:**
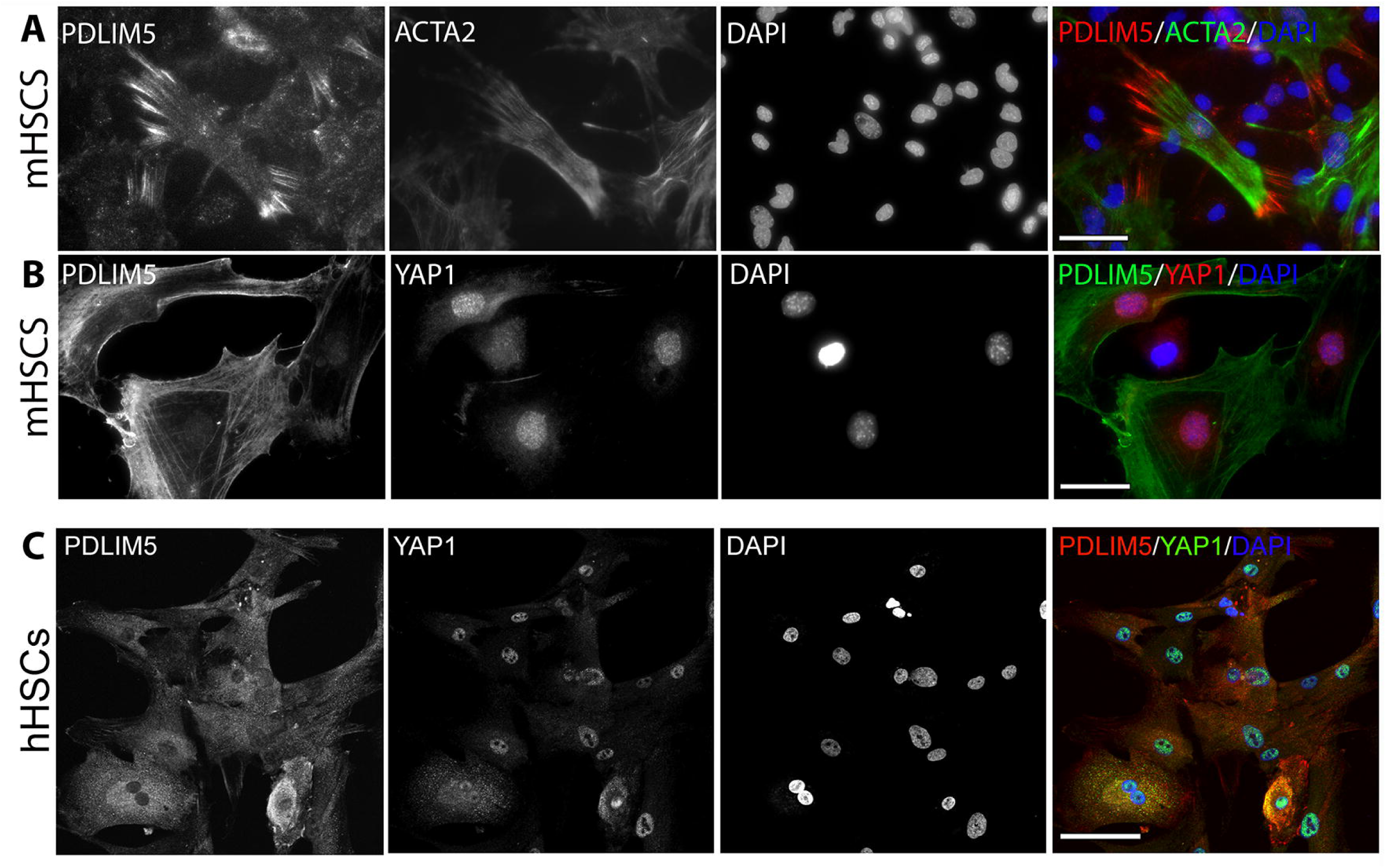
Dual immunofluorescence in primary HSCs. **A**. PDLIM5 (red) and ACTA2 (green); and **B**. PDLIM5 (green) and YAP1 (red), in mHSCs. **C.** PDLIM5 (red) and YAP1 (green) in primary human HSCs (hHSCs). DAPI nuclear stain (blue). Scale bars 100µm.

### PDLIM5 is involved in HSC Mechano-Sensing

PDLIM5 has been identified as a component of the integrin adhesome^4^ so we examined PDLIM5 localisation in HSCs relative to the focal adhesion protein paxillin during cell adhesion to fibronectin. After 90 minutes adhesion to fibronectin coated plastic LX-2 cells displayed (**Figure 5**) extensive spreading with PDLIM5 clearly detected in the cytoplasm and in punctate structures at the cell periphery. The pattern of staining is consistent with focal adhesions formed during cell adhesion and spreading. The merged image shows overlapping PDLIM5 (green) and paxillin (red) signals, particularly at the cell periphery in paxillin rich areas. In LX-2s PDLIM5 therefore has high spatial overlap with paxillin during fibronectin mediated adhesion, supporting a role for PDLIM5 in HSC adhesion and potentially in focal adhesion-associated signalling mechanisms during HSC activation.

**Figure 5:**
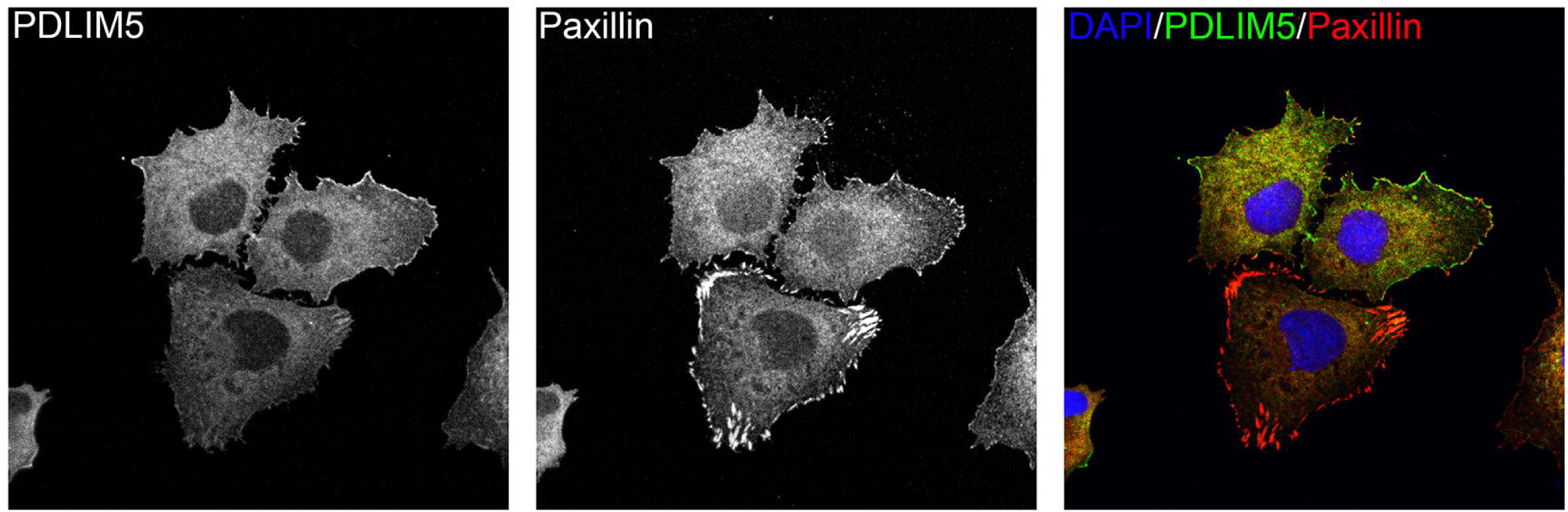
LX-2 cell adhesion on fibronectin coated plastic after 90minutes. PDLIM5 (green) and Paxillin (red).

In culture activated primary mHSCs, the myosin inhibitor blebbistatin has been shown to inhibit contraction and focal adhesion formation^30^. Blebbistatin did not alter *PDLIM5* expression in LX-2 cells (**Figure 6A**), but *ACTA2* expression was significantly (*p<0.01*) inhibited by blebbistatin both in the presence and absence of TGFb. This suggests *PDLIM5* expression is regulated independently of myosin contraction. Despite stable *PDLIM5* expression, the intracellular distribution of both PDLIM5 and YAP1 appeared altered following blebbistatin treatment (**Figure 6B**). Furthermore, we have previously observed that blebbistatin reduced nuclear size in LX-2 cells^27^, which was confirmed by current data (**supplemental figure 5**). These observations suggest that cytoskeletal tension may influence PDLIM5 function primarily through changes in localisation rather than transcriptional regulation.

**Figure 6:**
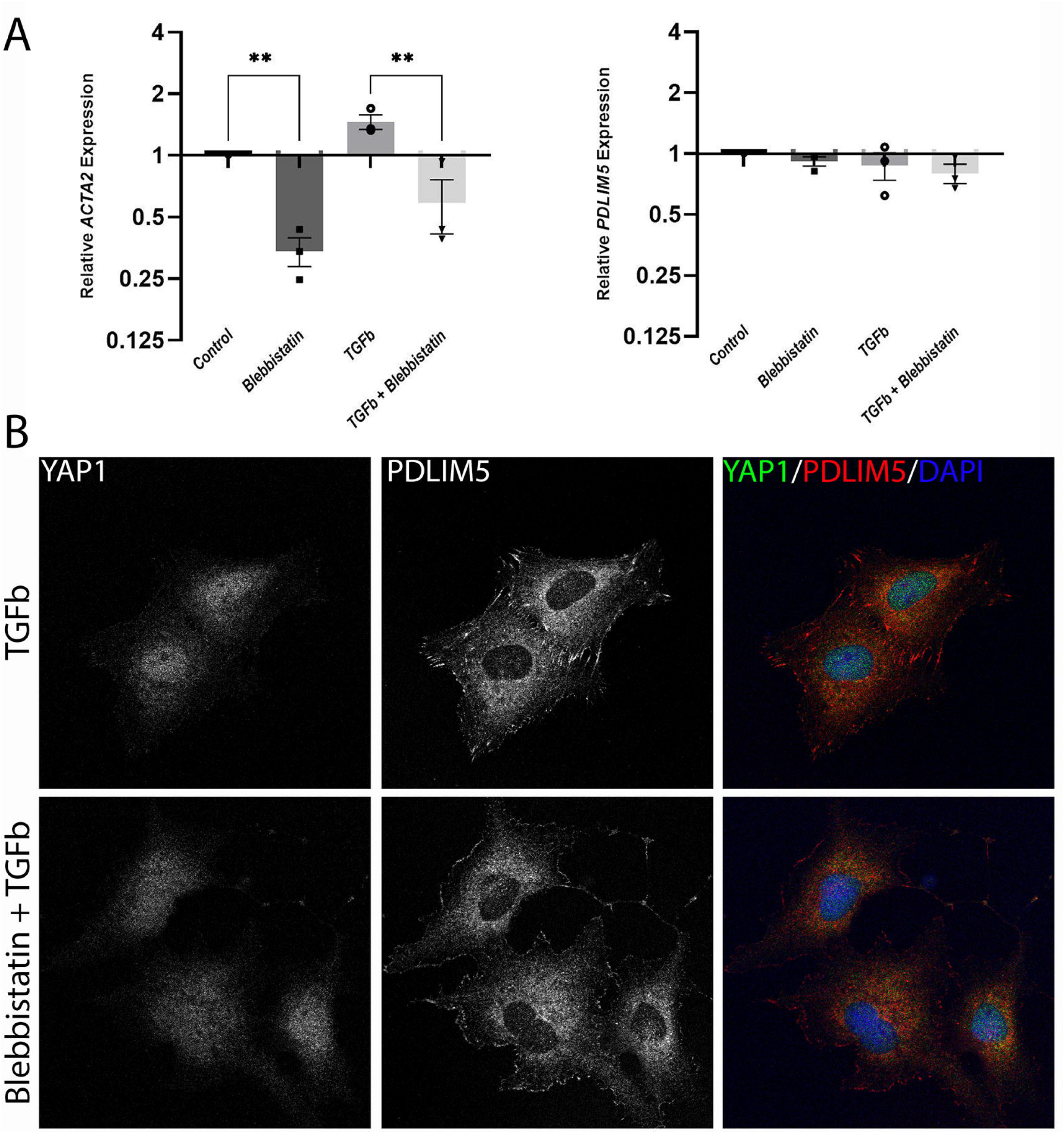
**A**. qPCR of LX-2 cells treated with 2 doses of 10µM blebbistatin at time zero and 6hours, +/- 5ng/µl TGFb. RNA was harvested at 24hrs. *ACTA2* expression was significantly inhibited in the presence of blebbistatin while *PDLIM5* was not affected (n=3, ** *p<0.01*). **B**. LX-2 cells were treated with 5ng/µl TGFb +/- 10µM blebbistatin and stained with antibodies against YAP1 (Green) and PDLIM5 (red) and nuclei labelled with DAPI (blue). Individual channels (grayscale) and merged images are shown.

Importantly PDLIM5 in primary mHSCs is associated with ACTA2 expression and cellular location, and PDLIM5 localisation changes in response to substrate stiffness (**Figure 7A**). In LX-2 cells YAP1 appeared to be more nuclear localised when cells were grown on stiff substrates (**Figure 7B**) in line with previous observations^8–10^. Furthermore, the distribution of PDLIM5 within the cytoplasm also appeared to change depending on substrate stiffness, with more PDLIM5 around the nucleus of cells grown on 12kPa substrates, as well as an apparent increase at the cell periphery (**Figure 7B**). To test the potential association of PDLIM5 with YAP1 in HSCs we performed co-immunoprecipitation and proximity ligation assays. Endogenous PDLIM5 was detected in YAP1 antibody pulldowns from LX-2 lysates (**Figure 8A, supplementary figures 10 and 11**). Proximity ligation assays revealed close spatial association between endogenous PDLIM5 and YAP1 within intact LX-2 cells (**Figure 8B**). Together, these findings support the existence of a PDLIM5-YAP1 signalling complex in HSCs that could be involved in modulating YAP1 mechano-sensing relevant to HSC activation.

**Figure 7.**
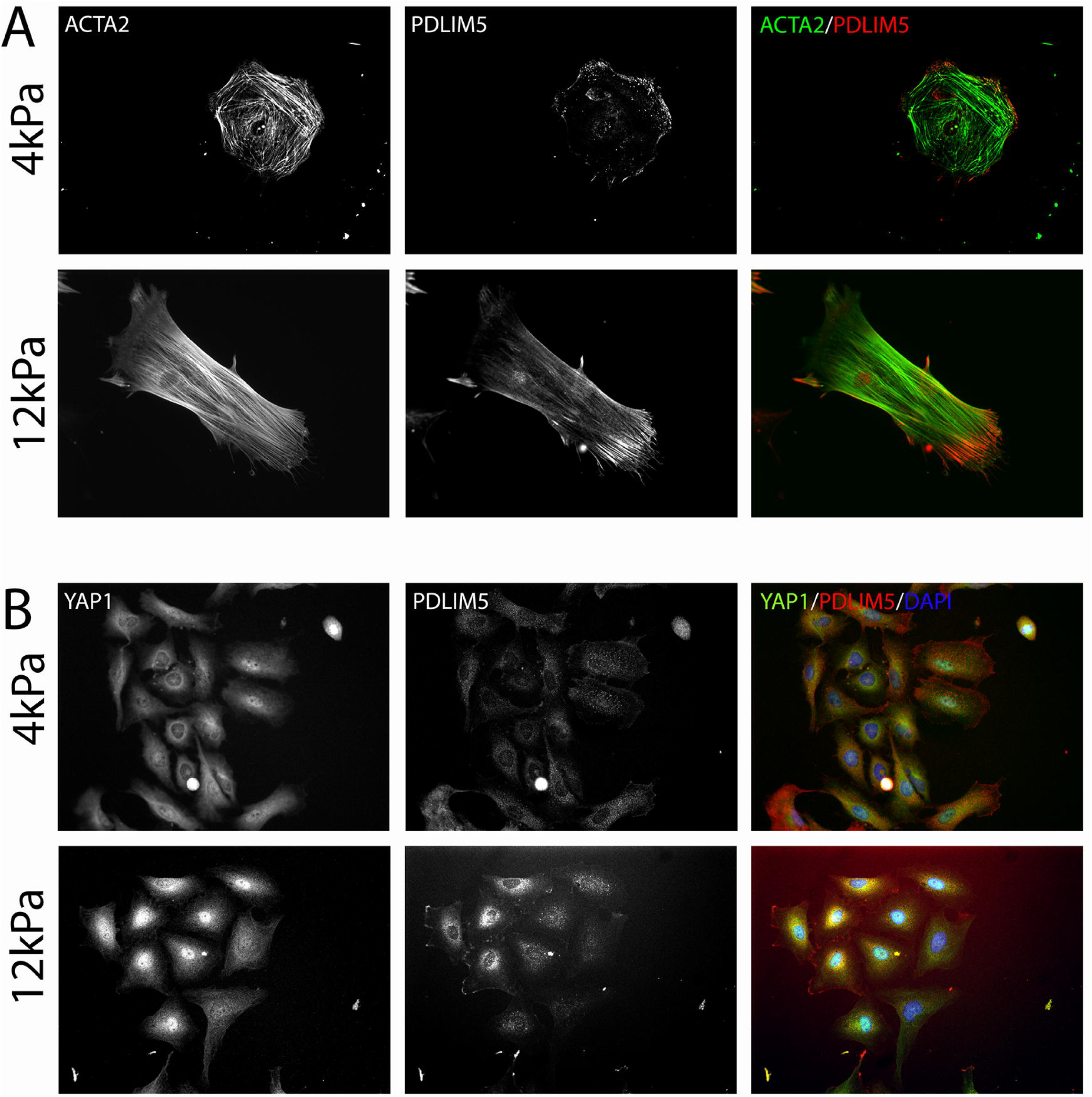
**A**. Primary mHSCs grown on 4kPa or 12kPa substrates stained for ACTA2 (green) and PDLIM5 (red). **B**. LX-2 cells grown on 4kPa or 12kPa substrates stained for YAP1 (green) and PDLIM5 (red).

**Figure 8:**
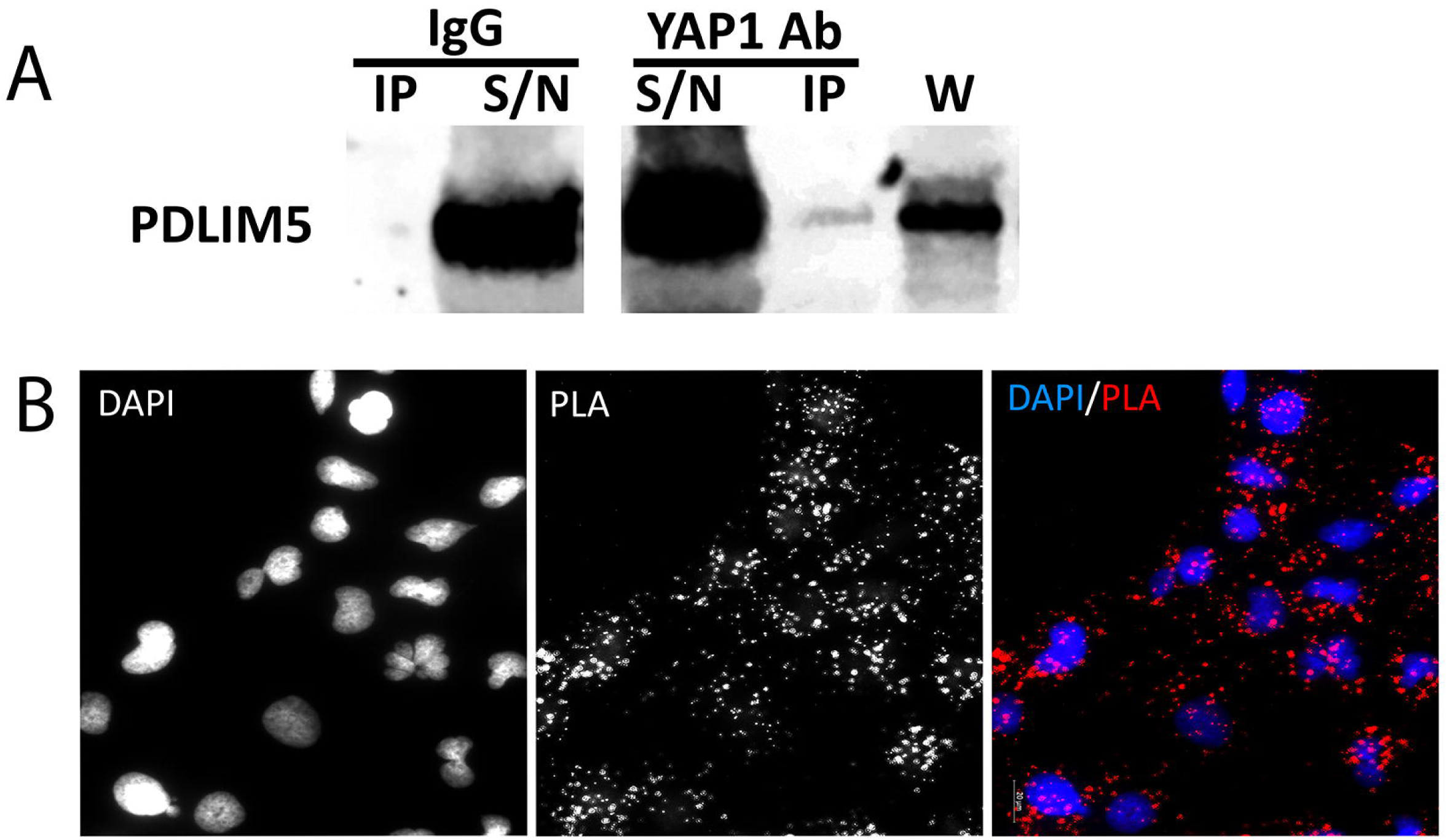
**A.** YAP1 antibody immunoprecipitation (IP) in LX-2 cells. PDLIM5 band is detected in the YAP1 IP. PDLIM5 was not detected in IPs with non-specific IgG (IP –immunoprecipitation, S/N – supernatant, W – whole cell lysate). **B**. Proximity ligation assay (PLA, red) with YAP1 and PDLIM5 antibodies in LX-2 cells. DAPI nuclear stain (blue).

### Targeting PDLIM5 inhibits fibrotic gene expression and mechano-signalling

To interrogate the function of PDLIM5 we used 3 parallel approaches, pharmacological inhibition, siRNA, and CRISPR interference (CRISPRi). Low dose (50nM) paclitaxel has been shown to inhibit PDLIM5-mediated cell signalling^31^ IC_50_ values for cell viability of over 350nM into the micromolar range are reported in NSCLC cell lines treated with paclitaxel for 24 hours^32^. In LX-2 cells 50nM paclitaxel induced altered cell morphology (**Figure 9A**) and 25nM was sufficient to reduce the expression of HSC activation-associated genes including *ACTA2 (p<0.05)*, *COL1A1* and the pro-fibrotic YAP1 target *CTGF*^22^ *(p<0.01)* (**Figure 9B**). In addition, paclitaxel reduced collagen I protein expression in LX-2s and inhibited the upregulation of collagen I protein expression in response to TGFb (**Figure 9C**). This effect on collagen I is in agreement with previous observations made in HSCs, using far higher concentrations (200nM) of paclitaxel^33^. Importantly, YAP1 localisation appears to be less nuclear in LX-2s treated with 50nM paclitaxel (**Supplementary figure 6**). These observations support the hypothesis that PDLIM5 is involved in HSC activation in a mechanism potentially involving the regulation of YAP1 localisation.

**Figure 9.**
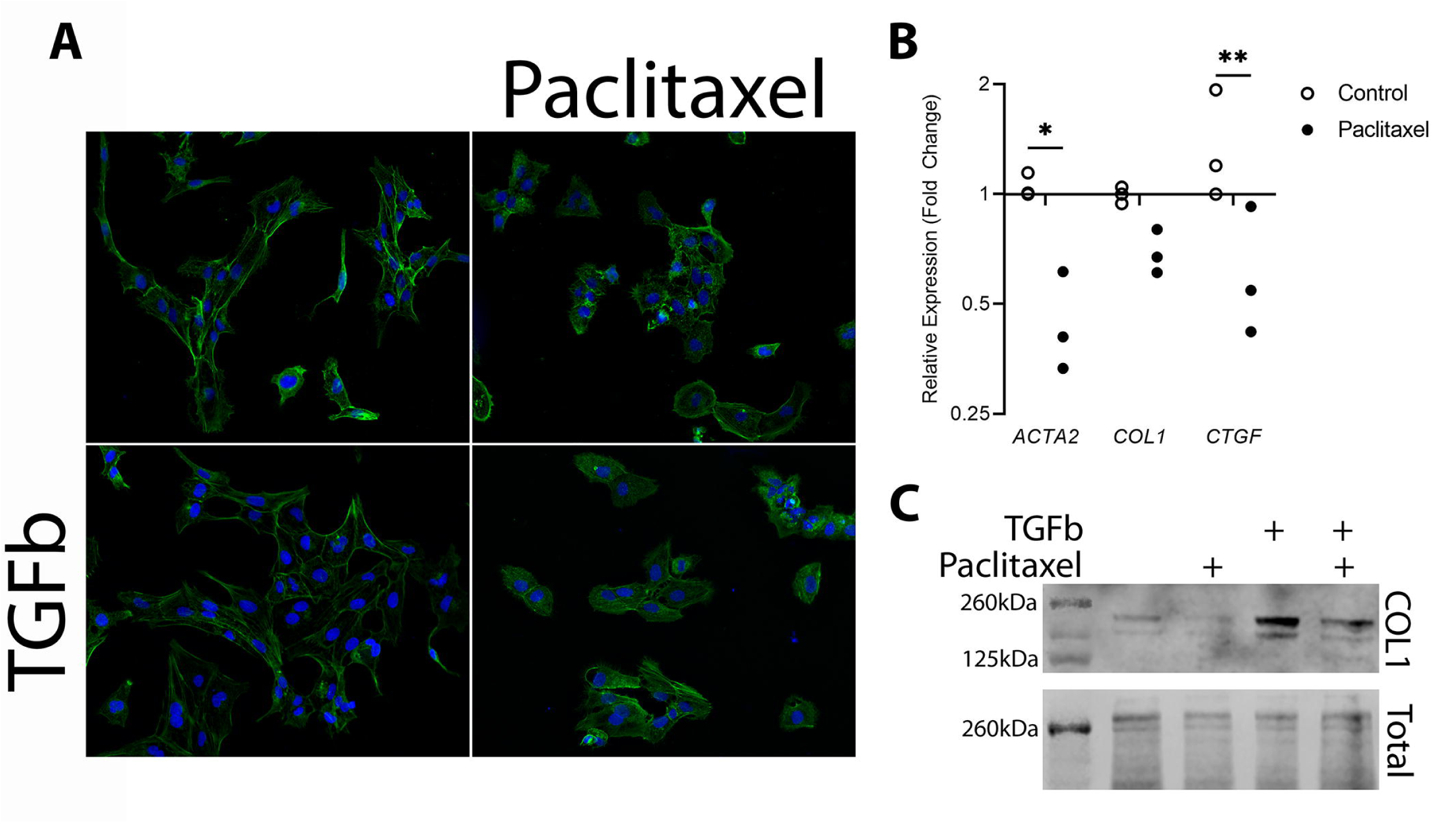
Paclitaxel as a tool for pharmacological inhibition of PDLIM5. **A.** Paclitaxel (50nM) treatment of LX-2 cells +/- TGFb: Phalloidin (green) and DAPI (blue). **B**. *ACTA2*, *COL1* and *CTGF* qPCR in LX-2 cells treated with 25nM paclitaxel *(* p<0.05*, *\*\* p<0.01*). **C**. Western blot of COL1 in lysates from LX2 cells treated with TGFb and/or 50nM Paclitaxel.

To further investigate the function of PLDIM5 in HSCs a pool of 4 siRNAs targeting PDLIM5 was used. In agreement with the paclitaxel data *PDLIM5* siRNA significantly reduced *ACTA2* and *COL1* expression in LX-2 cells (*p<0.001)*, and expression of the pro-fibrotic YAP-1 target *CTGF* was also reduced (*p<0.01*, **Figure 10A**). Reduction of PDLIM5 protein *(p<0.001*) was confirmed by western blotting (**Figure 10B and C**) and was accompanied by significantly (*p<0.01*) reduced ACTA2 expression (**Figure 10B and C**). Importantly, when PDLIM5 expression was reduced, nuclear localisation of YAP1 also appeared to be reduced (**Figure 10D**). These observations suggest that PDLIM5 may contribute to maintaining HSCs in an activated phenotype, possibly as part of a mechano- sensitive mechanism involving the control of YAP1 localisation.

**Figure 10.**
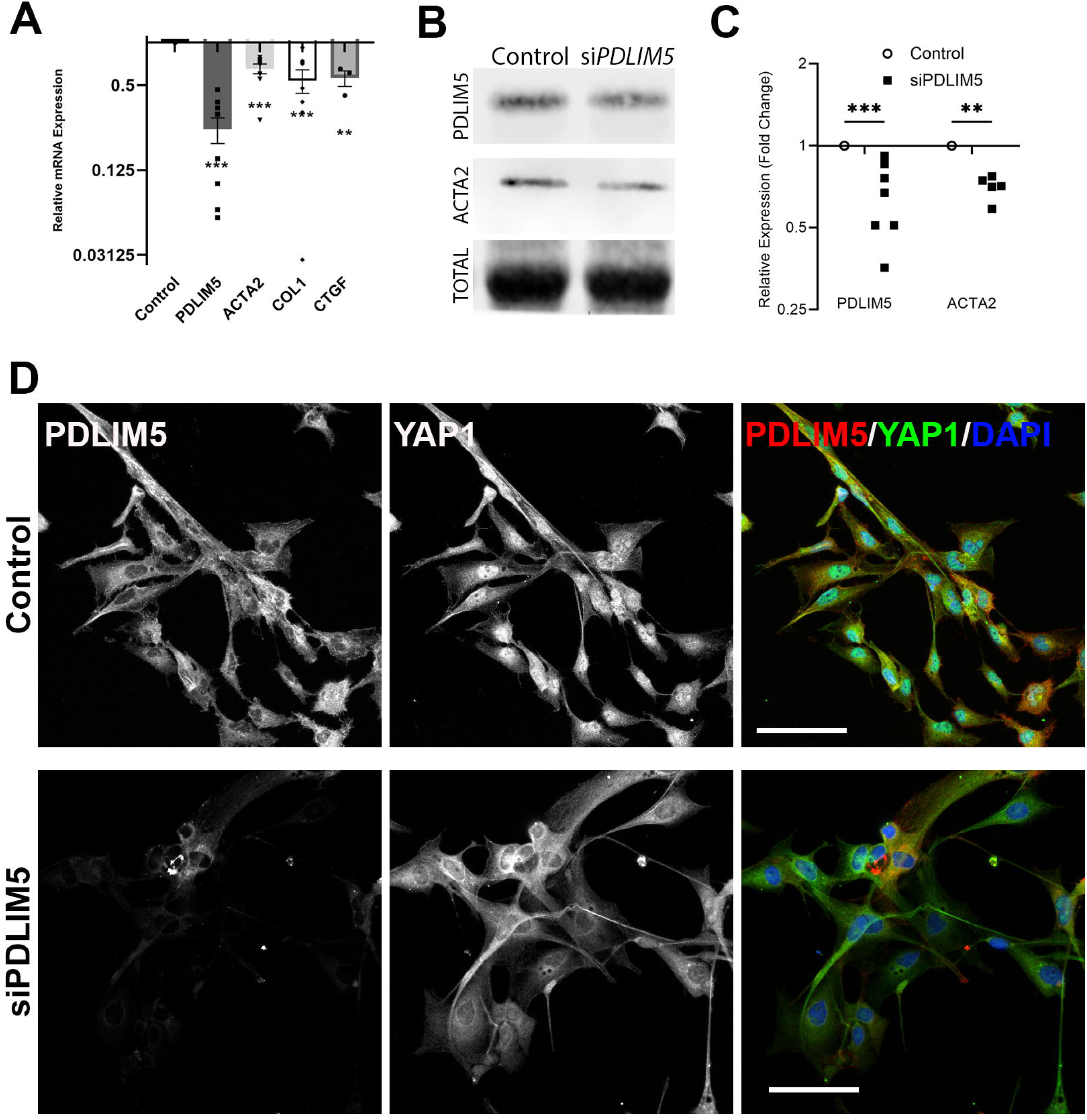
Inhibition of *PDLIM5* expression using siRNA in LX-2 cells reduces fibrotic gene expression and inhibits nuclear translocation of YAP1. **A**. qPCR comparing gene expression in control and si*PLDIM5* LX-2 cells. Mean relative expression (fold change) on log2 scale, +/-SEM with replicates plotted (*n=5 or 3 for CTGF, *** p<0.001, ** p<0.01*) using *GusB* and *RPLO* as reference genes. **B.** Western blot showing PDLIM5 and ACTA2 protein expression in control and *PDLIM5* siRNA LX-2 cells. Total is total protein stain. **C**. Densitometry of PDLIM5 and ACTA2 plotted as relative protein expression. (*n=5, *** p<0.001, ** p<0.01*). **D**. YAP1 (green) and PDLIM5 (green) in control and si*PDLIM5* LX-2 cells. Scale bars are 100µm.

To validate the siRNA data, *PDLIM5* expression was also targeted by using CRISPRi^34^ to repress *PDLIM5* transcription. In further support of a role for *PDLIM5* in the control of HSC phenotype, and consistent with the data from paclitaxel and siRNA experiments, inhibiting *PDLIM5* expression by CRISPRi significantly reduced *ACTA2* expression (**Figure 11**).

**Figure 11:**
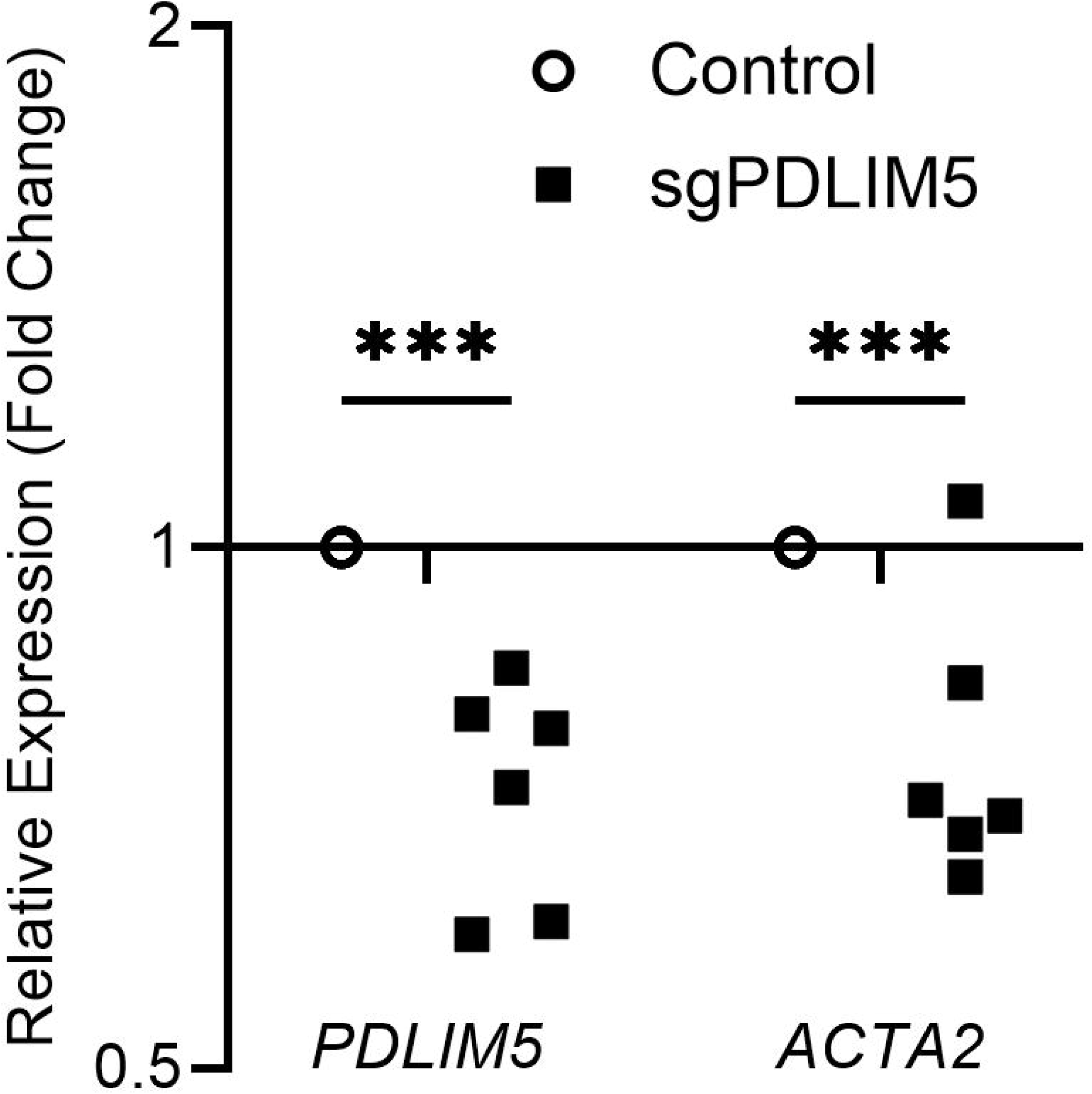
CRISPRi of *PDLIM5* in LX-2 cells caused reduced *ACTA2* expression. Reference genes *HPRT* and *RPLP0*, n=6, *\*\*\* p<0.001*).

## DISCUSSION

Mechano-signalling via YAP1 is an established driver of HSC activation and liver fibrosis^8–10^, but the mechanisms that control YAP1 activity in HSCs remain poorly defined. This data shows for the first time that the adhesome component PDLIM5^4^ is present in activated HSCs *in vivo* in human and in experimental mouse liver fibrosis. We show that PDLIM5 localisation in cells within regions of scarring, and within cells co-labelled for α-Sma *in vivo* is consistent with a role for PDLIM5 in cytoskeletal and adhesion-associated signalling in HSCs during fibrosis. PDLIM5 associates with YAP1 and inhibition of PDLIM5 reduces YAP1 nuclear localisation and pro-fibrotic gene expression. Together these observations suggest that PDLIM5 may contribute to mechanosensitive HSC activation via YAP1.

Mechano-signalling is a key regulator of HSC activation and fibrosis progression^15,18–20^. PDLIM5 is an established component of the cellular adhesome^4^ and therefore a plausible mediator of force-dependent signalling, including via YAP1^26^. In HSCs this is further supported by the observed localisation of PDLIM5 with stress fibres and focal adhesion associated proteins such as paxillin, as well as by changes in PDLIM5 distribution in response to changes in substrate stiffness.

YAP1 drives mechano-activation of HSCs^8,10^. Co-IP and PLA data now support proximity between endogenous PDLIM5 and YAP1 in HSCs. Reduced nuclear localisation of YAP1 in the context of reduced *PDLIM5* expression indicates PDLIM5 regulates YAP1 activation or localisation in HSCs. This is consistent with previous reports supporting a role for PDLIM5 in YAP1 mechano-regulation in epithelial cells^26^.

Using multiple approaches to target PDLIM5 we observed reduced expression of *ACTA2*, *COL1A1* and *CTGF*. siRNA, CRISPRi and pharmacological approaches all support a functional role for PDLIM5 in maintaining the activated HSC phenotype. This regulation of HSC phenotype may occur in part via YAP1. The mechanism by which PDLIM5 influences YAP1 signalling in HSCs remains to be defined. It seems reasonable to suggest that PDLIM5 acts as a scaffold linking the actin cytoskeleton and adhesion complexes to YAP1-regulation. Alternatively changes in the cytoskeleton and cellular tension due to PDLIM5 inhibition may indirectly alter YAP1 localisation. Further investigation of the PDLIM5 interactome, adhesion signalling and YAP1 phosphorylation and localisation in HSCs are needed to better establish the likely mechanism.

Our experiments to explore the role of PDLIM5 in HSC activation were based on observing PDLIM5 protein in HSCs during fibrosis *in vivo*. However, many of our functional experiments were done using LX-2 cells. Future investigations should extend to include primary HSCs, and ideally *in vivo* models of liver fibrosis or liver organoids^35^. It also remains to be firmly established whether there is a direct interaction between PDLIM5 and YAP1, although this hypothesis is supported by both Co-IP and PLA. In addition to siRNA and CRISPRi we have used low dose paclitaxel as a pharmacological tool to inhibit PDLIM5^31^, but the effects of paclitaxel in HSCs may not be limited to acting via PDLIM5.

In summary, these data identify PDLIM5 as a previously overlooked component of the HSC adhesome with a likely role in HSC activation and mechano-sensing. Further investigation of PDLIM5 function in HSCs is necessary to understand how PDLIM5 supports HSC mechano-signalling via YAP1.

## METHODS

### Human Tissue

Human liver was collected with informed consent and ethical approval (National Research Ethics Service, REC 14/NW1260/22) from the Manchester Foundation Trust Biobank.

### CCl4 *In Vivo* Fibrosis Model

C57Bl/6J Mice were housed, maintained, and experiments performed in accordance with UK Government Home Office regulations and with the approval of the University of Manchester Ethical Review Committee. Liver fibrosis was induced by 2μl intraperitoneal (i.p.) injections of sterile CCl4 (Sigma, UK) per g body weight in a ratio of 1:3 by volume in olive oil, or olive oil alone (control) twice weekly for 8 weeks. Age matched 8–12-week-old male mice were used. Liver tissue was fixed in 4% PFA and processed for histology as previously reported^8^.

### Primary Cell Isolation and Cell Culture

Primary mouse HSCs were harvested under terminal anaesthesia by perfusion with pronase and collagenase via the portal vein. HSCs were isolated from the resulting cell suspension with an Optiprep (Axis-Shield Diagnostics Ltd) density gradient as previously^8,36,37^. Animals were housed and maintained, and cell isolations performed under approval from the University of Manchester Ethical Review Committee and in accordance with UK Government Home Office regulations.

Primary human HSCs were isolated from liver tissue obtained with informed consent and ethical approval (National Research Ethics Service, REC 14/NW1260/22) from the Manchester Foundation Trust Biobank. Liver tissue was finely dissected and incubated at 37□°C for 40minutes in digestion buffer (0.5□mg/ml collagenase B, 0.2□mg/ml pronase), and HSCs were isolated from the resulting cell suspension using Optiprep density gradients (Axis-Shield Diagnostics Ltd) as previously described^27,38^.

Activation of primary HSCs was achieved by 7 days of culture on cell culture plastic^37–39^. Isolated primary HSCs were characterised for expected changes in gene expression and phenotype^8,37^. HSCs were cultured in high glucose DMEM enriched with Na-Pyruvate, L-glutamine, pen-strep and 16% serum.

The LX-2 immortalised male human HSC line was established by Professor Scott Friedman (Mount Sinai School of Medicine)^40^. LX-2 cells were maintained in high glucose DMEM enriched with Na-pyruvate, L-glutamine, pen-strep and 1% serum and routinely screened for mycoplasma.

TGFb was used at a final concentration of 5ng/µl. Blebbistatin was used at 10µM. Paclitaxel was used at 25nm or 50nm as stated.

### siRNA Knockdown

*PDLIM5* was targeted for knockdown with SMARTpool ON-TARGET *plus* siRNA (Horizon Discovery, L-006930-00-0005 Lot 190725). SMARTpool uses a pool of 4 siRNA for increased potency and specificity. Target sequences were 1. AAGAAUAGGCGAUGUGGUU; 2, CAACAUGCCUCUCACAAUC; 3. AGAAUAAGAUUAAGGGUUG; and 4. GAAUAUGACUCUGCAAAGA. LX-2 cells were transfected with siRNA using Dharmafect Transfection Reagent 1 (Horizon Discovery). A pool of 4 Non-targeting siRNAs (Horizon Discovery, ON-TARGETplus Non-targeting Control Pool, D-001810-10) was used as the control for siRNA experiments. LX-2 cells were transfected with siRNA using Dharmafect 1.

### dCas9 LX-2 Cell Lines for CRISPR Interference/Activation

To establish dCas9-CRISPRi cell lines, LX-2 cells were transduced with hCMV CRISPRi dCas9-SALL1-SDS3 lentiviral particles (Horizon Discovery). For dCas9-CRISPRa cells were transduced with hMCV-Blast-dCas9-VPR (Horizon Discovery). Stable lines were established by Blasticidin (5ug/ml) selection.

dCas9 cells were transfected (Dharmafect, Horizon Discovery) with pooled guide RNA targeting *PDLIM5* for either CRISPRi or CRISPRa (Horizon Discovery, CRISPRi SMARTpool sgRNA *CF-006930-01-0005*; CRISPRa SMARTpool sgRNA *PG-006930-01- 0005*) using Dharmafect 1. Non-targeting gRNA was used as control (Horizon Discovery SMARTpool sgRNA).

### qPCR

RNA was extracted using RNEasy purification kits (Qiagen). RNA was converted to cDNA using high-capacity RNA-to-cDNA kits (Life Technologies). Relative transcript abundance was calculated using ddCt in SYBR Green based qPCR assays. When comparing quiescent and activated mHSCs at least 3 biological repeats were used. i.e. Using HSCs prepared from at least 3 different mouse livers. Reference genes were selected using CFX Maestro (BioRad). For mHSC qPCR reference genes were *GusB* and *RPLO* in combination. For LX-2 experiments reference genes were *GusB* and *ActinB* unless indicated otherwise in figure legends. LX-2 experiments used at least 3 replicates. Intron spanning primers were designed using NCBI Primer Blast^41^. Primer sequences are provided in supplemental table 1.

### Western Blot

Protein lysates were collected by removing all cell culture media, washing in PBS and lysing cells in RIPA buffer (Sigma) with protease inhibitors (Sigma). Protein concentrations were measured by standard BCA assay (Biorad). For western blot proteins were denatured using Laemlli buffer and separated using standard acrylamide gel electrophoresis. Proteins were transferred to nitrocellulose, blocked in casein and used for antibody labelling. Antibodies used were ACTA2 (Abcam, Ab7817), COL1 (Southern Biotech, 1310-01), and PDLIM5 (Sigma Prestige Antibodies, HPA016740). Total protein was detected using Revert Stain (Li-cor Biosciences). Primary antibody binding was visualised using IRDYe secondary antibodies (Li-cor Biosciences) and membranes were scanned using Li-Cor Odyssey imaging XF system. Densitometry analysis was performed using Fiji/ImageJ^42^. Digital blot images were converted to 8-bit grayscale, and rectangular regions of interest of identical size were applied to each lane. Band intensities were quantified using the Gel Analysis tool. Total protein was used for signal normalisation. See supplemental data for original western blot images.

### Immunoprecipitation

Immunoprecipitation was carried out as previously described^8^. Whole cell lysates were incubated overnight at 4°C with either normal IgG (Cell Signalling Technology) or YAP1 antibody (Santa-Cruz, sc-271134) and anti-IgG beads (Protein G beads, Cell Signalling Technology). Original western blot images are provided in the supplementary data.

### Histology and immunohistochemistry

Liver tissue was fixed in 4% PFA and processed for histology as previously reported^9,37,43^. Immunohistochemistry for PDLIM5 (Sigma Prestige Antibodies, HPA016740, 1:100) was detected using imPRESS anti-Rabbit IgG Peroxidase (Vector Labs) and diaminobenzidine (DAB) and counterstained with toluidine blue.

### Immunofluorescence

Primary HSCs or LX-2 cells were plated onto chamber slides or fibronectin coated hydrogels of the indicated stiffness (Softwell plates, Cell Guidance Systems). Cells were washed twice in PBS and fixed for 10mins in 4% PFA in PBS. Following 2 PBS washes cells were stored at 4°C in PBS until staining. Cells and tissue were stained with Alexa Fluor 488 Phalloidin (Invitrogen, 1:1000) and/or primary and secondary antibodies (Alexa Fluor, Invitrogen): ACTA2 (Leica Biosystems, SMA-R-7-CE, 1:100)^8^, Paxillin (BD Bioscience, 610052, 1:500), PDLIM5 (Sigma Prestige Antibodies, HPA016740, 1:100), YAP1 (Santa Cruz, Sc-101199, 1:200)^8^. Liver tissue sections were treated with TrueVIEW Autofluorescence quenching kit (Vector Labs). For analysis of immunofluorescence images Fiji/ImageJ^42^ was used. Image processing and analysis parameters are provided in the Supplementary Methods. Individual measurements were averaged per field of view and subsequently averaged within each biological replicate to avoid pseudoreplication.

### Proximity Ligation Assay

Protein-protein interactions were detected using Duolink Proximity Ligation Assay (Sigma). Antibodies used were YAP1 (Santa Cruz, sc-271134) and PDLIM5 (Sigma Prestige Antibodies, HPA016740).

### Statistical Analysis

For qPCR data analysis Bio-Rad CFX Maestro was used. Expression data (2^-ΔΔCt^) was exported to GraphPad Prism 9 for ANOVA with Sidak post hoc test. Data are presented as fold change relative to control on a log scale with a baseline of 1.

Data from immunofluorescence images is presented as replicate means derived from at least 3 fields of view per condition and analysed in GraphPad Prism 9 using unpaired two-tailed t- tests.

## Author Contributions

JP and JH conceived and designed experiments. KPH, KW, DA, and VSA contributed to experimental design. JP, KW, DA, LI, EFJ, HK, JMM, LB, AM, RP and EJ performed experiments. JP, JH and KW analysed the data. JP, JH, TM and KPH wrote and contributed to editing the manuscript.

