## Supplemental Data for "PDLIM5 Modulates YAP1 Localisation and Fibrogenic Gene Expression in Hepatic Stellate Cells"

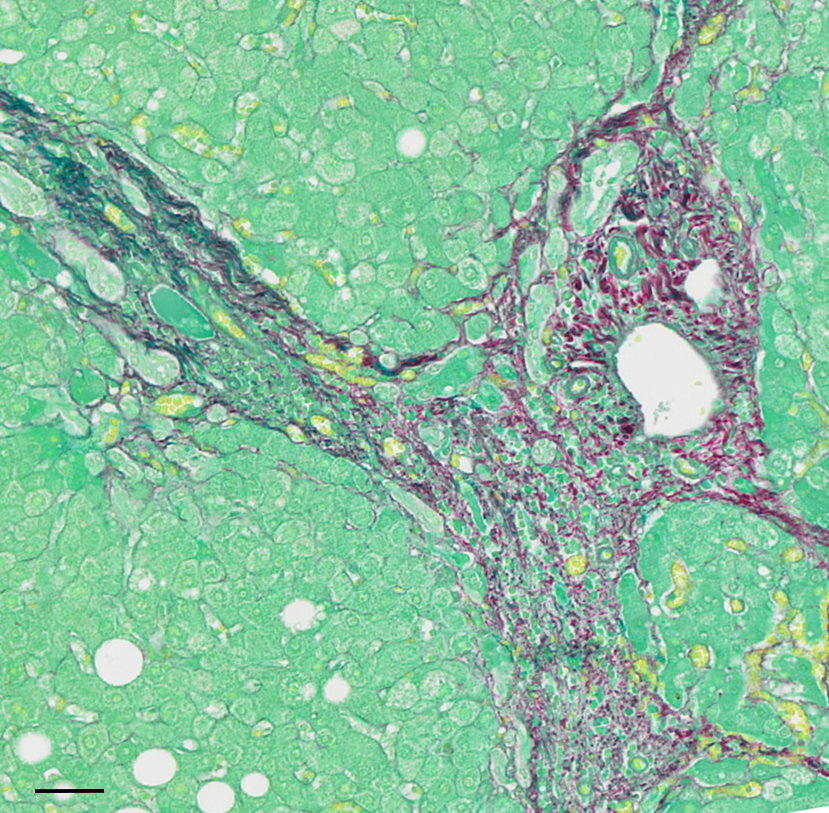
**Supplementary Figure 1:** Representative portal region from a near-serial section of human liver corresponding to figure 1A, stained with Sirius red. Scale bar is 50um.

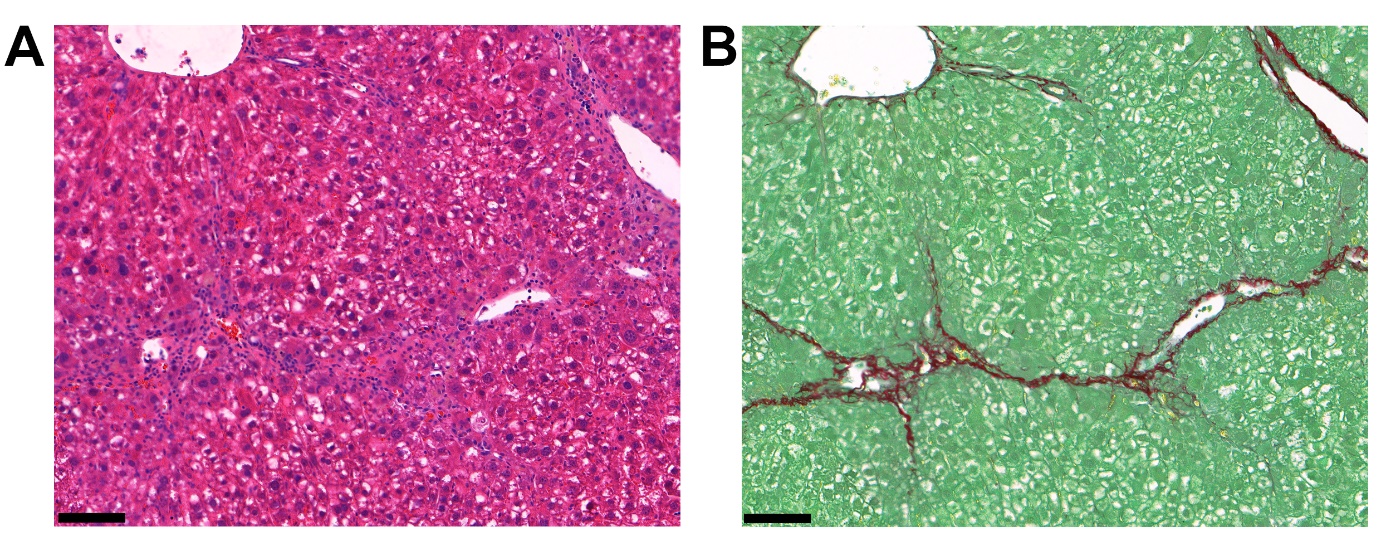
**Supplemental Figure 2:** Representative region from near-serial sections of mouse liver corresponding to figure 1B stained with haematoxylin and eosin (A) and Sirius red (B). Scale bars are 100um.

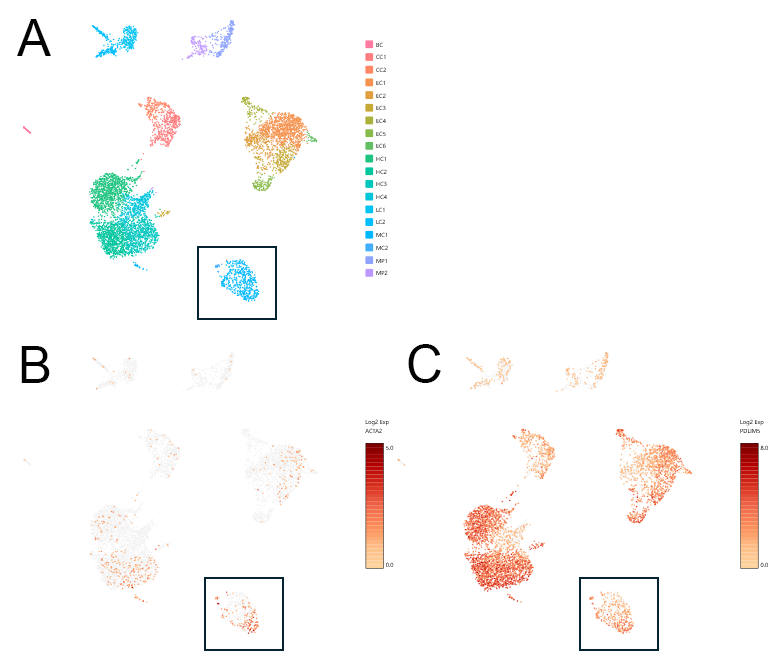
**Supplemental Figure 3:** Uniform Manifold Approximation and Projection (UMAP) plot of human liver single nucleus RNA sequencing (snRNA-Seq) data from Hammond et al^1^. Box indicates mesenchymal cells (A). Expression levels of *ACTA2* (B) and *PDLIM5* (C) demonstrate a population of *ACTA2* positive mesenchymal cells express *PDLIM5*.

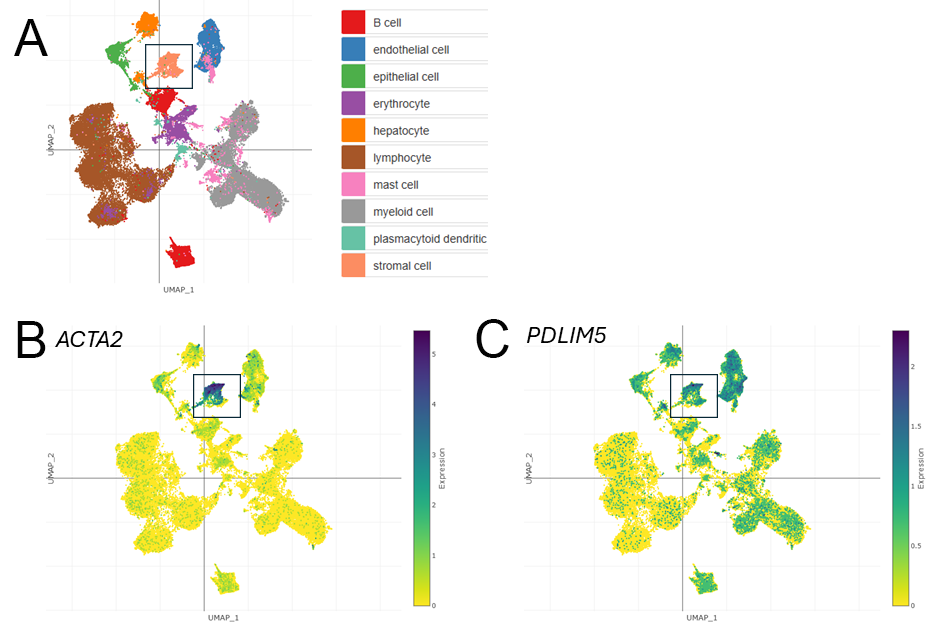

**Supplemental Figure 4**: UMAP plot of human liver scRNA-seq data from Fabre et al^2^. The stromal cell population (A, box) contains *ACTA2* (B) positive cells that also express *PDLIM5* (C). Data accessed via Broad Institute Single Cell Portal, <https://singlecell.broadinstitute.org/single_cell>

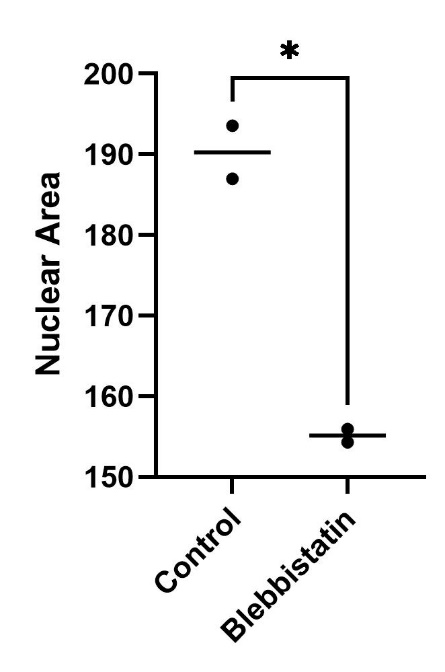

**Supplemental Figure 5**: Nuclear size in LX-2 cells treated with Blebbistatin (*n=2, p<0.05*)

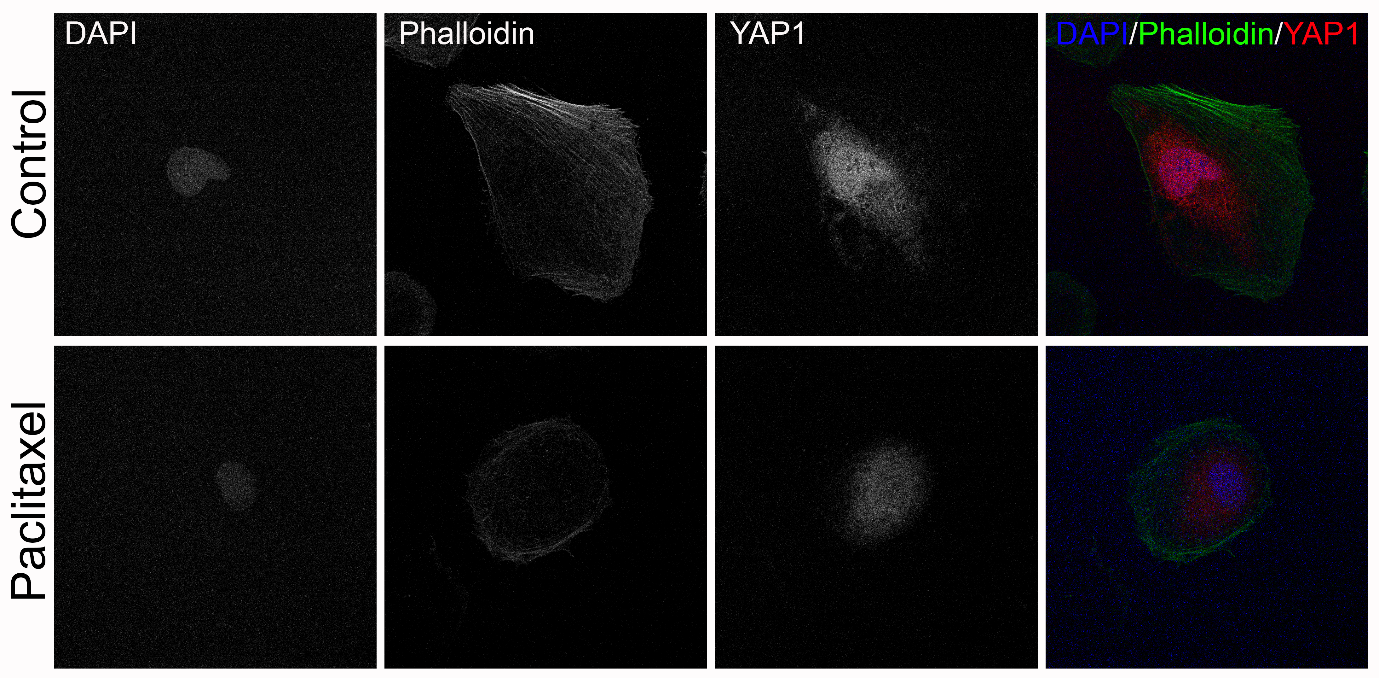
**Supplemental Figure 6:** LX-2 cells treated with 5ng/ul TGFb +/- 50nM paclitaxel and labelled with DAPI (blue), phalloidin (green) and YAP1 (red).

**Supplementary Figure 7**: Original western blots of LX-2 cell lysates probed with COL1 and PDLIM5 antibodies used to make composite in figure 3C. M, marker.

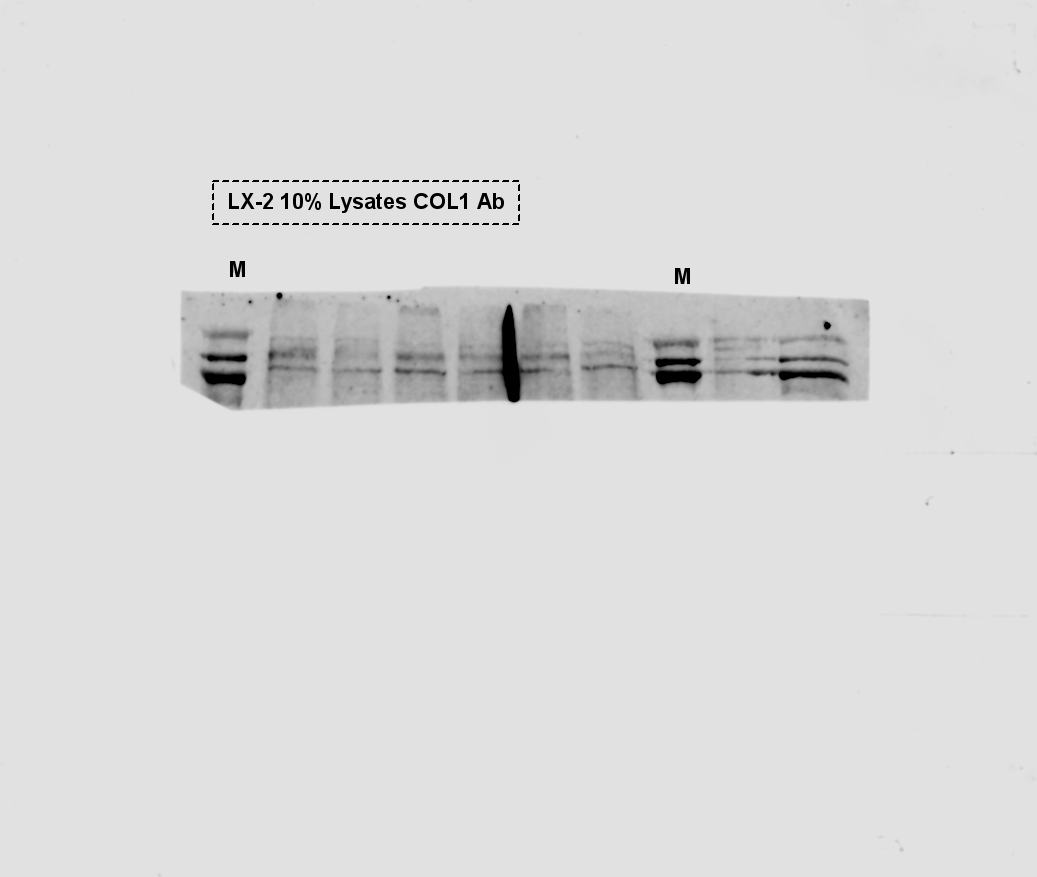

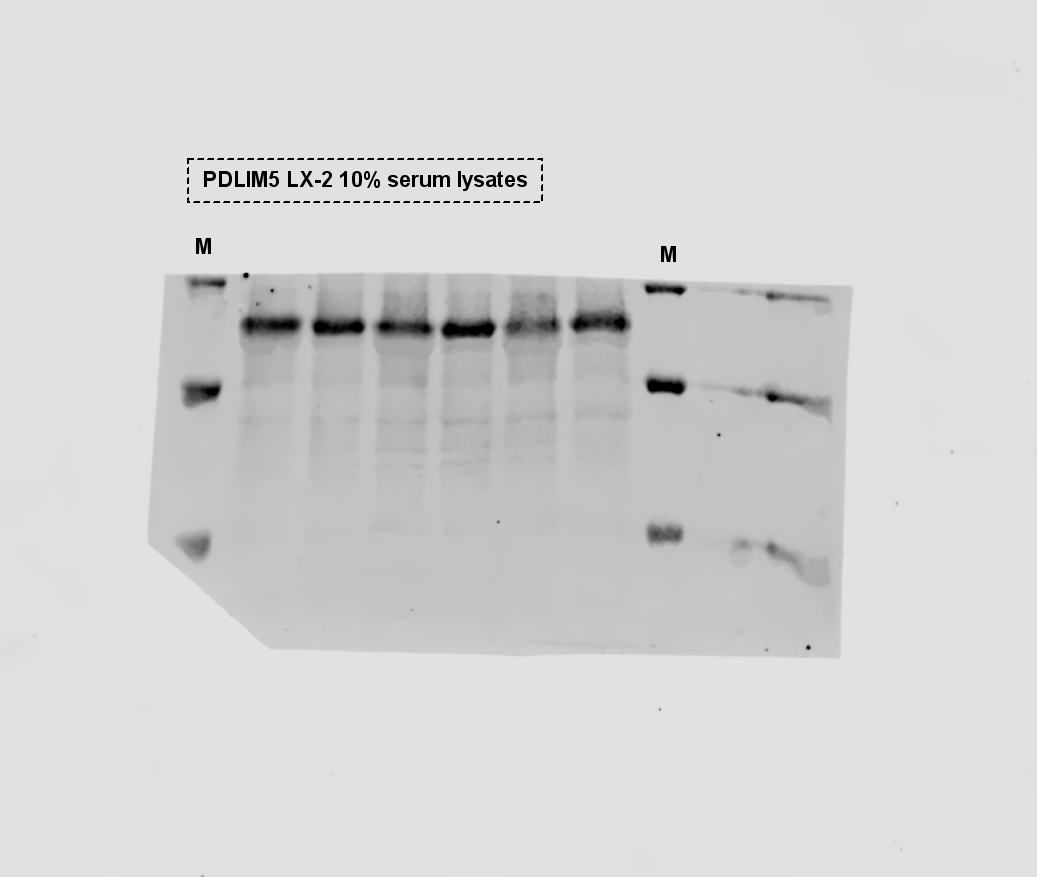

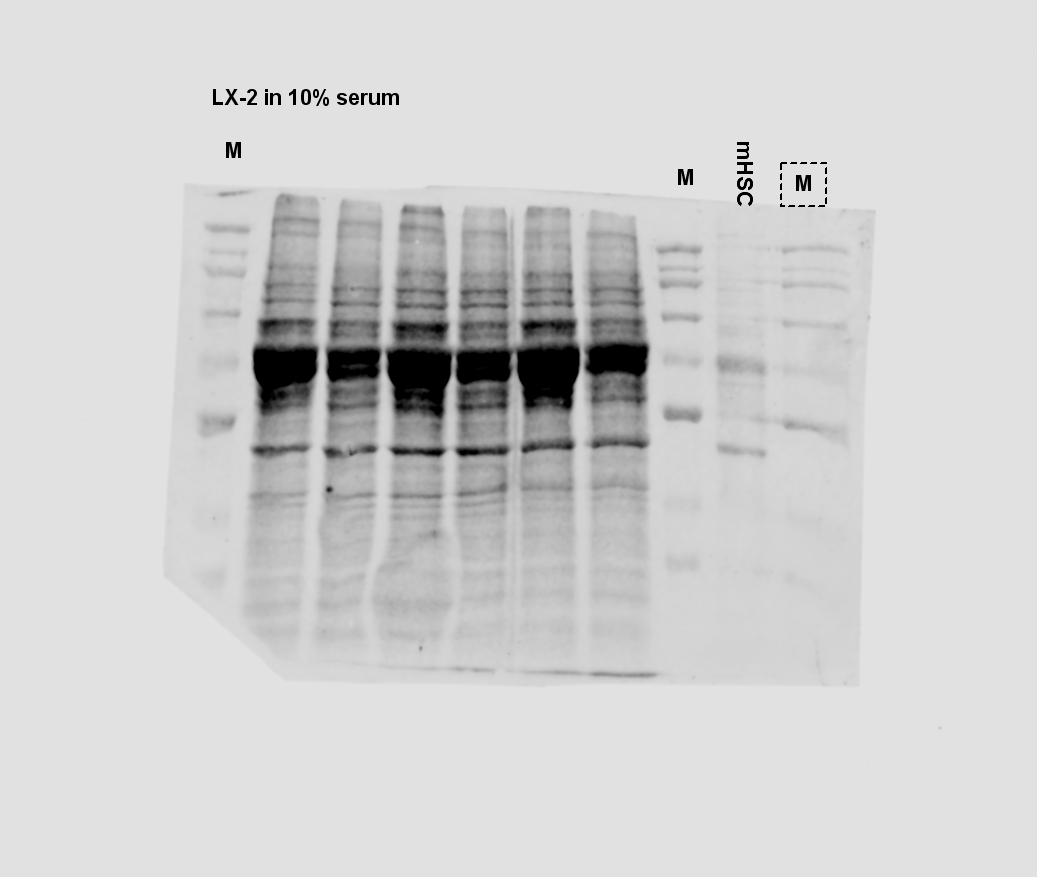

**Supplementary Figure 8**: Original western blot of LX-2 cell lysate probed with A-SMA antibody. Used to make composite in figure 3C.

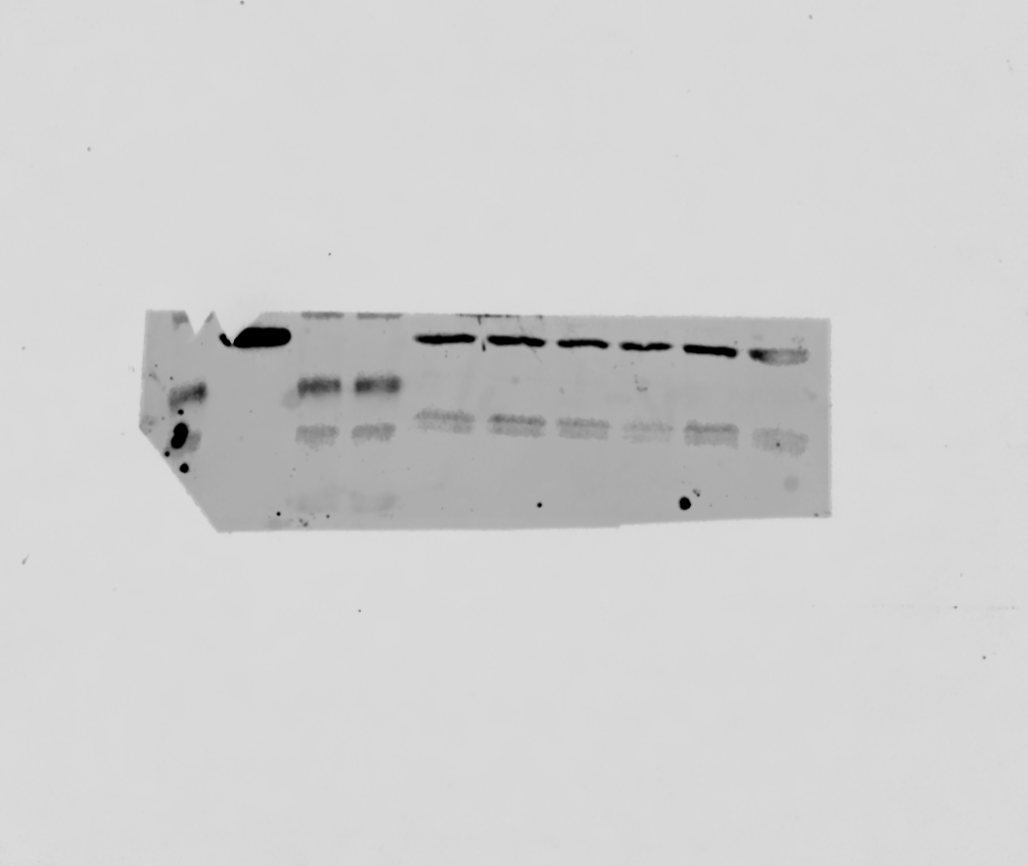

**Supplementary Figure 9:** Original western blot of primary mHSC lysates probed with A-SMA (green) and PDLIM5 (red) antibodies. Used to make composite in figure 3D. Q, quiescent, A, culture activated. M, marker.

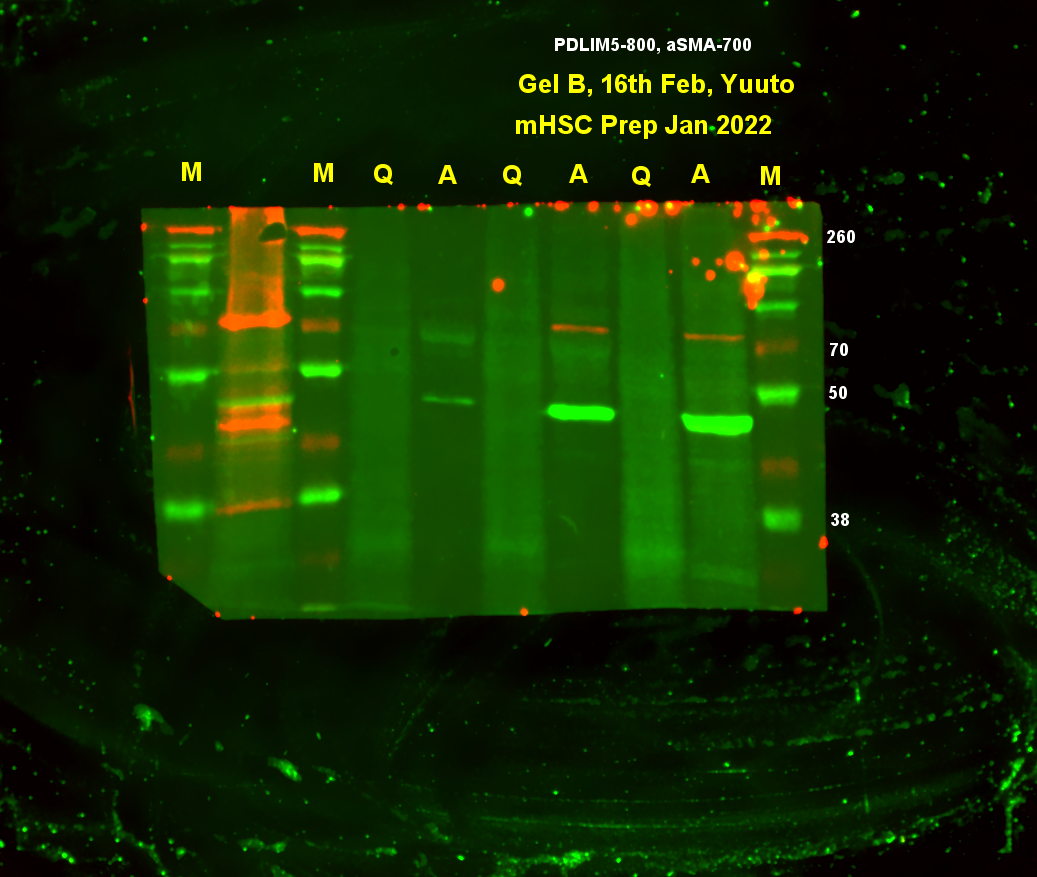

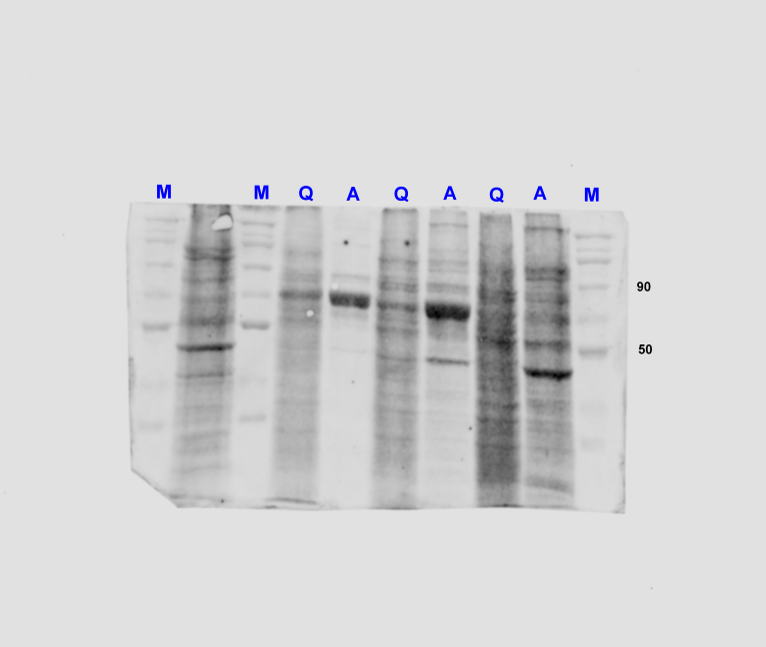

**Supplementary Figure 10:** Original blot of YAP-1 immuno-precipitates probed for PDLIM5. Used in figure 8.

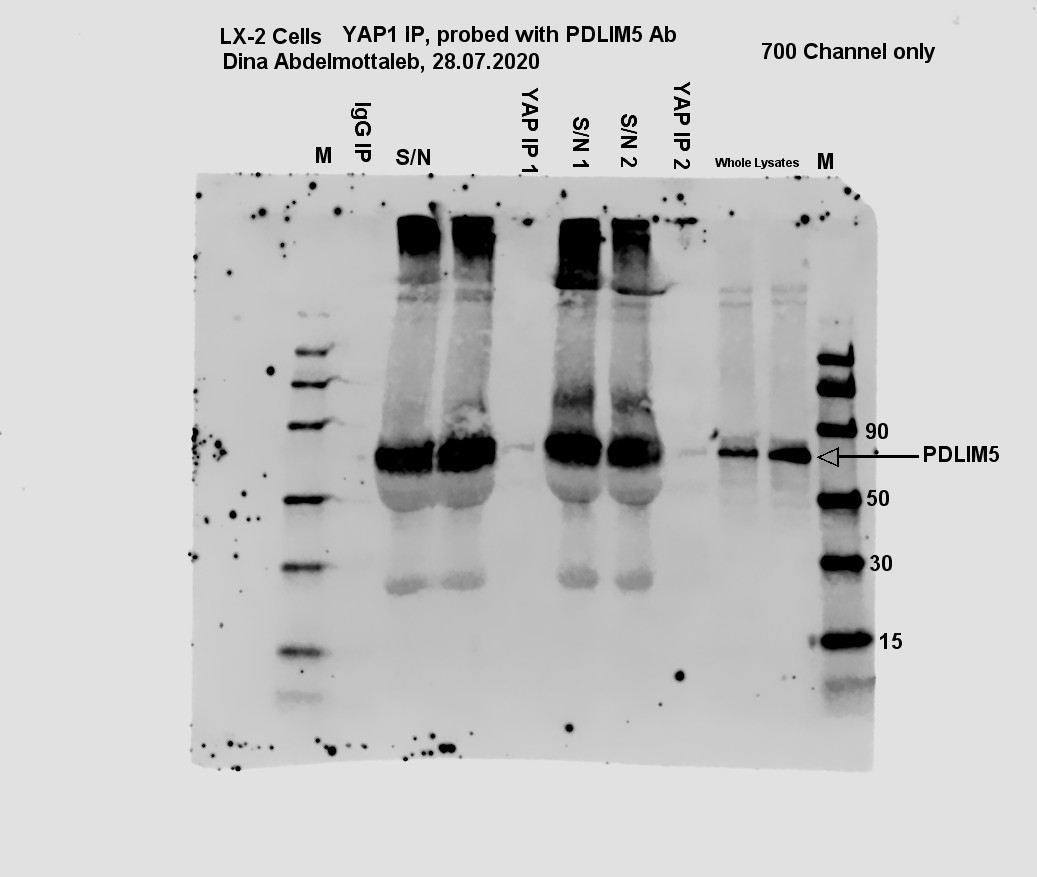

**Supplementary Figure 11:** Original blot of YAP-1 immuno-precipitates stripped and probed with YAP-1 antibody.

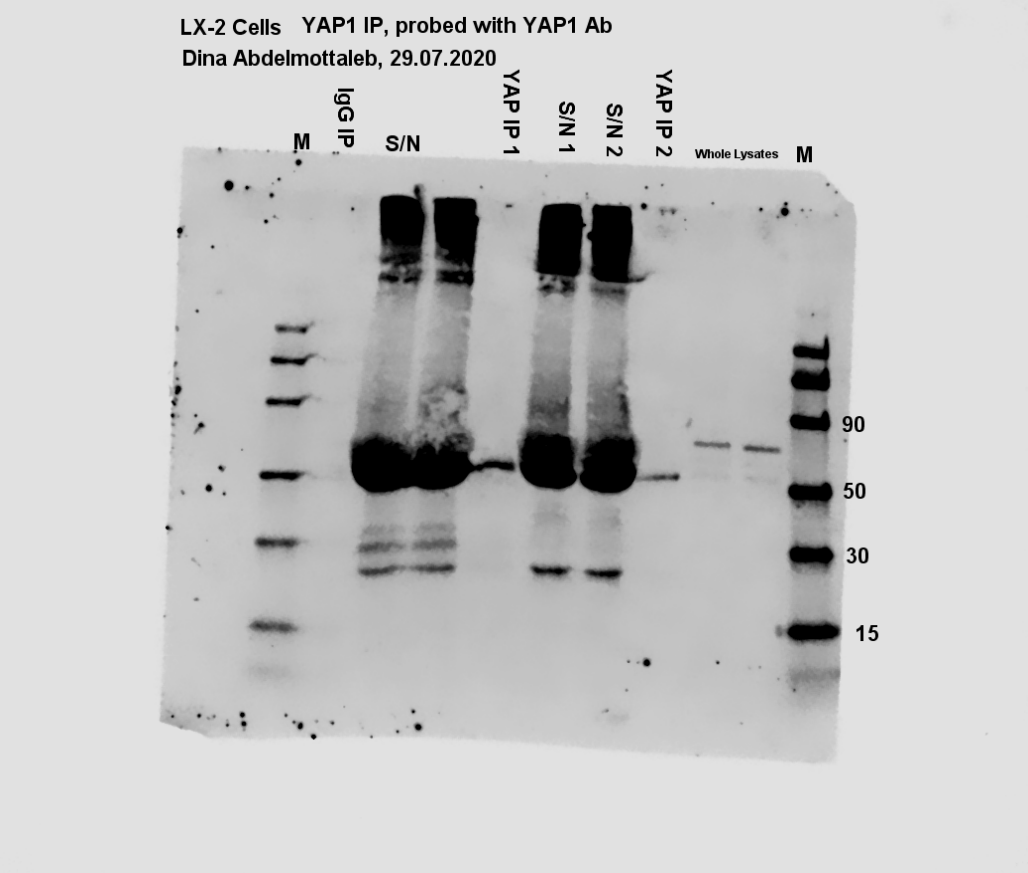

**Supplementary Figure 12:** Original western blot image of LX-2 lysates treated with paclitaxel (Taxol) and probed with COL1 antibody. Used in figure 9C.

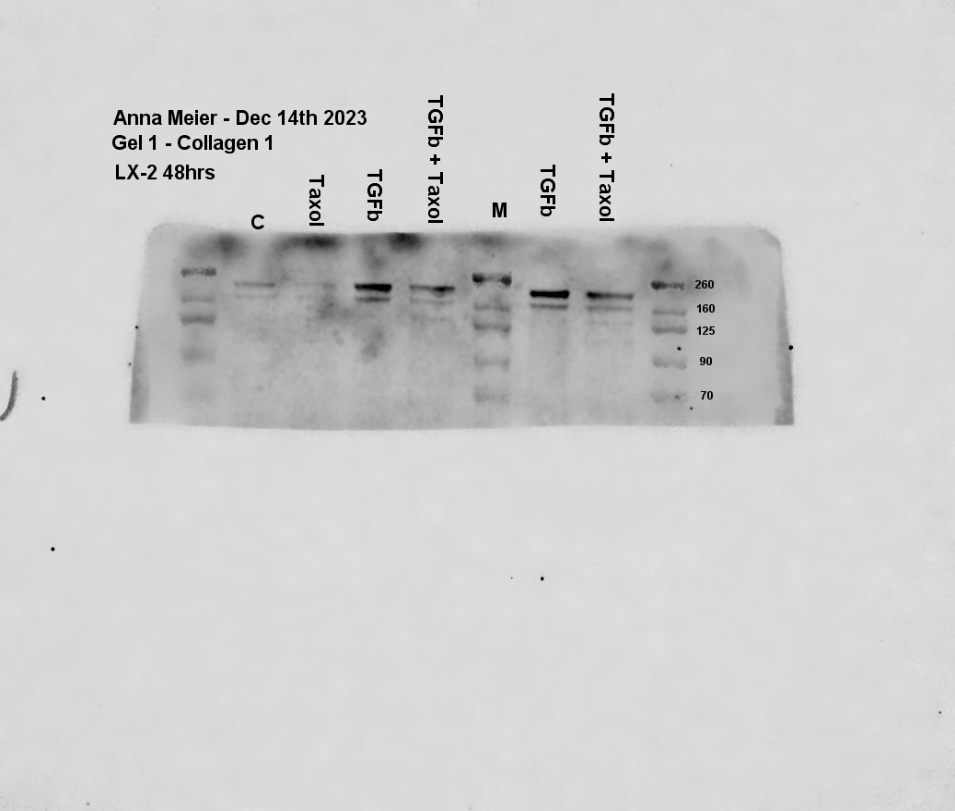

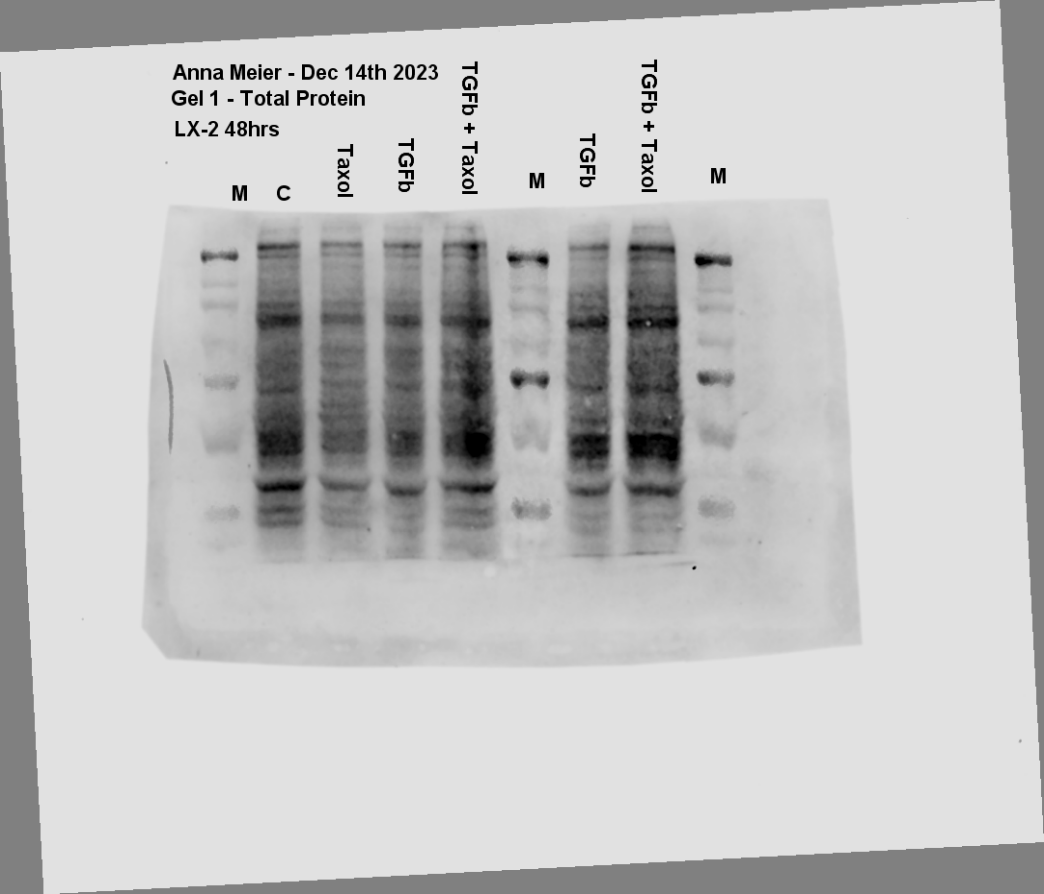

**Supplementary Figure 13:** Original western blots of LX-2 si*PDLIM5* lysates probed with a-SMA and PDLIM5 antibodies. These blots were used for densitometry, and the boxed regions are shown in Figure 10C with lane order reversed to show control (scrambled. S2) first.

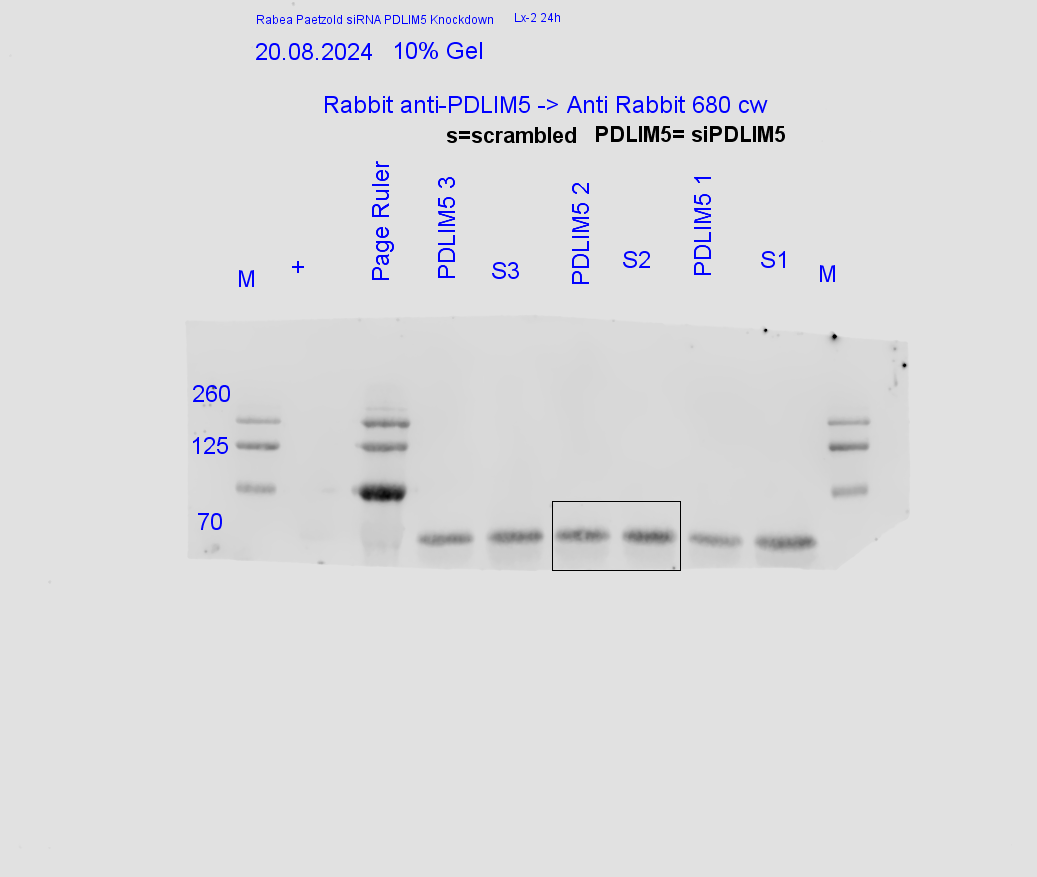

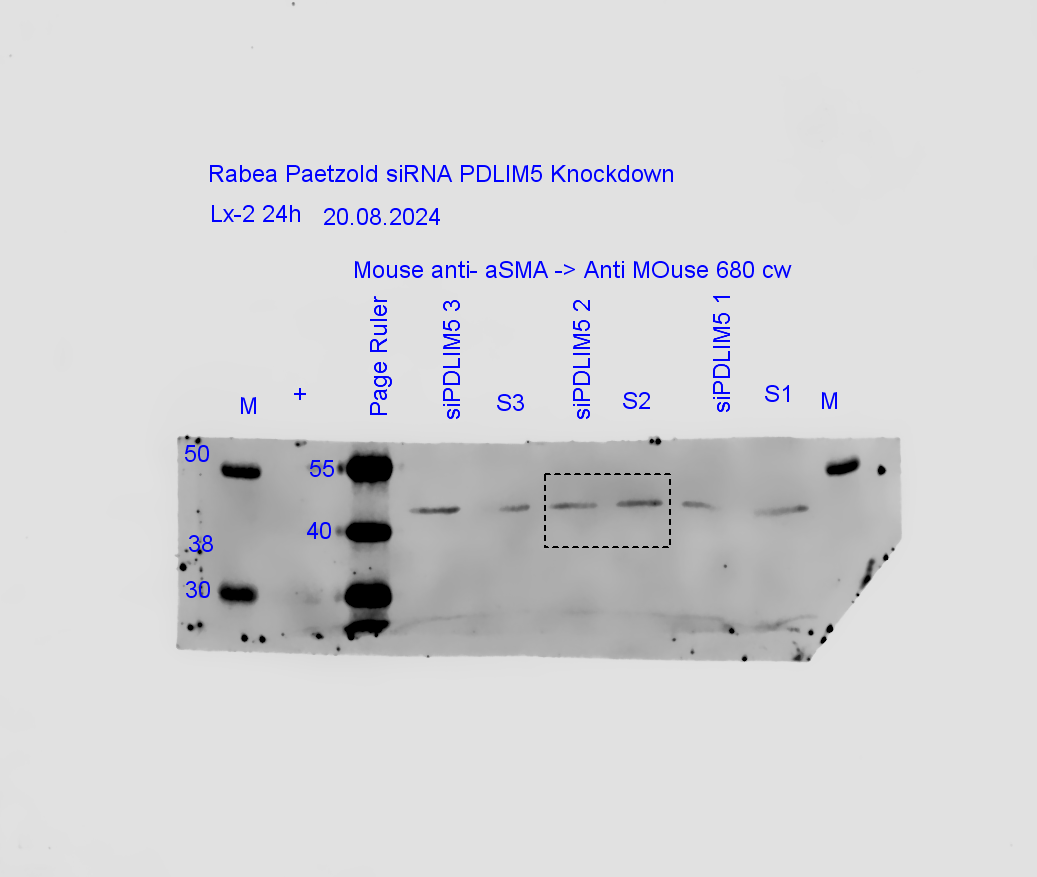

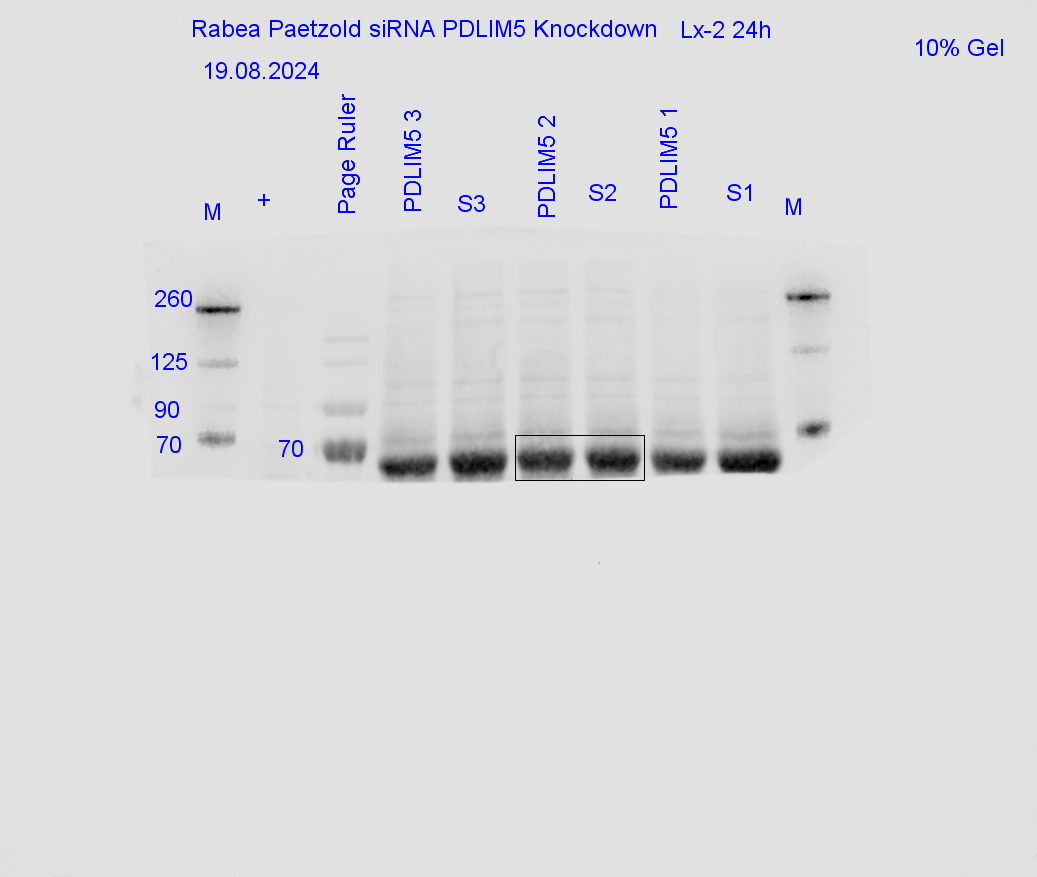

**Supplementary Figure 12**: Original western blots of LX-2 si*PDLIM5* lysates probed with a-SMA and PDLIM5. These blots were used with those in supplemental figure 8 for the densitometry data in Figure 10.

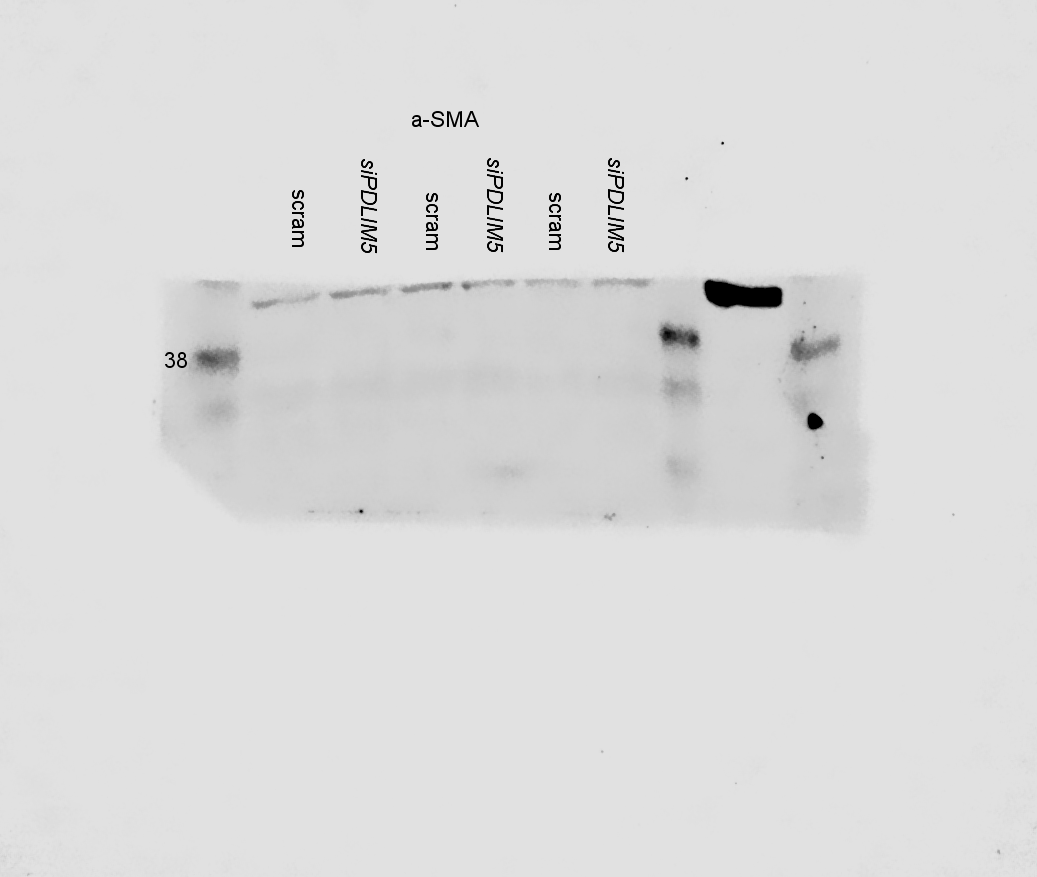

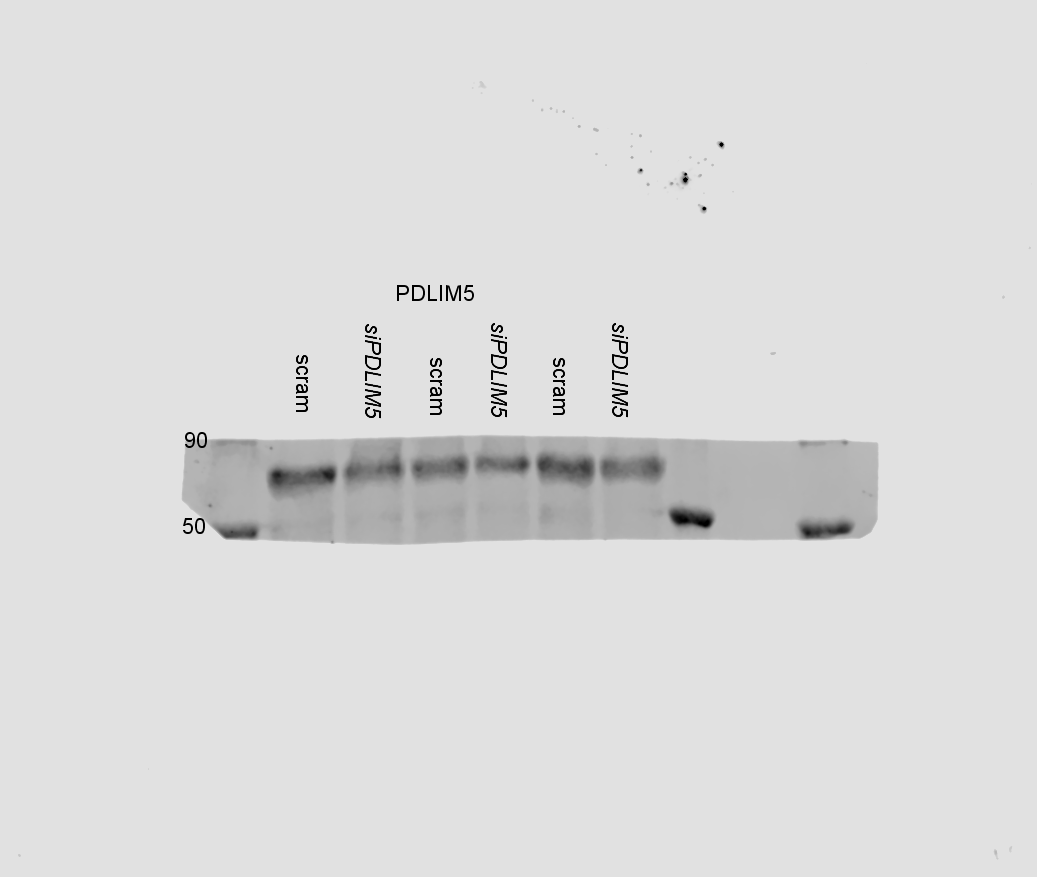

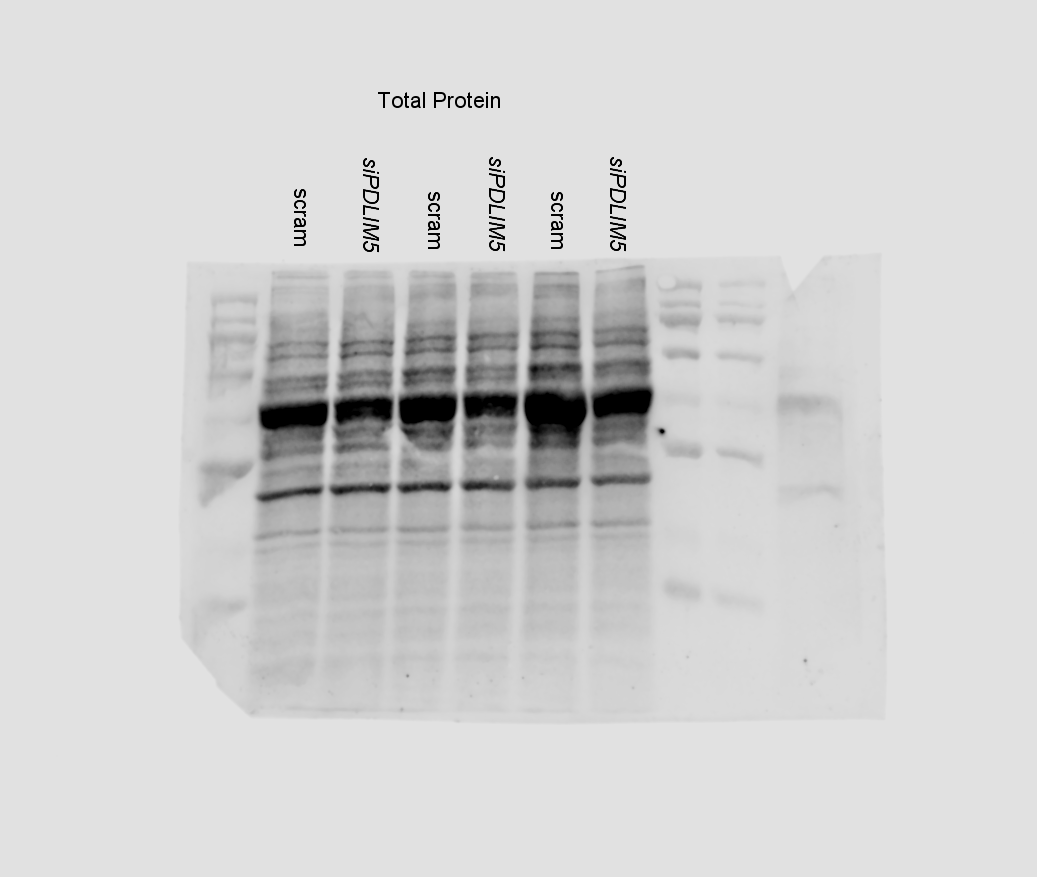

1	Hammond, N. L. *et al.* Spatial gene regulatory networks driving cell state transitions during human liver disease. *EMBO Molecular Medicine* **17**, 1452-1474 (2025). <https://doi.org:10.1038/s44321-025-00230-6>

2	Fabre, T. *et al.* Identification of a broadly fibrogenic macrophage subset induced by type 3 inflammation. *Science Immunology* **8**, eadd8945 (2023). <https://doi.org:doi:10.1126/sciimmunol.add8945>
