## Supplemental Methods for "PDLIM5 Modulates YAP1 Localisation and Fibrogenic Gene Expression in Hepatic Stellate Cells"

**Table 1.** Primer sequences.

| **Target Gene** | **Species** | **NCBI Accession** | **Forward Primer** | **Reverse Primer** |
| --- | --- | --- | --- | --- |
| *GusB* | Mouse | NM_010368.2 | TGGCTGGGTGTGGTATGAAC | TCCCATTCACCCACACAACT |
| *RPLP0* | Mouse | NM_007475.5 | CGTCCTCGTTGGAGTGACAT | TAGTTGGACTTCCAGGTCGC |
| *ACTA2* | Mouse | NM_007392.3 | GCCATCTTTCATTGGGATGGA | CCCCTGACAGGACGTTGTTA |
| *CTGF* | Mouse | NM_010217.2 | AGAACTGTGTACGGAGCGTG | GTGCACCATCTTTGGCAGTG |
| *PDLIM5* | Mouse | NM_019809.3 | TTCCGTCCAGAAGGGTGAAC | AAGGGCCGTGGTGCTTTATT |
| *GusB* | Human | NM_000181 | CTCATTTGGAATTTTGCCGATT | CCGAGTGAAGATCCCCTTTTT |
| *RPLP0* | Human | NM_001002.4 | CGTCCTCGTGGAAGTGACAT | TAGTTGGACTTCCAGGTCGC |
| *ActinB* | Human | X00351.1 | ccaaccgcgagaagatga | ccagaggcgtacagggatag |
| *ACTA2* | Human | Knerr at al^1^ and Goldberg et al^2^. | CCGACCGAATGCAGAAGGA | ACAGAGTATTTGCGCTCCGAA |
| *COL1* | Human | NM_000088.3 | TGTTCAGCTTTGTGGACCTCCG | CGCAGGTGATTGGTGGGATGTCT |
| *PDLIM5* | Human | NM_006457 | GGGTTGTACAGGCTCTTTGA | CCGTGGTGCCTTATTGTAGG |
| *CTGF* | Human | NM_001901.3 | CATCTTCGGTGGTACGGTGT | TTCCAGTCGGTAAGCCGC |

**Table 2**. Nuclear Area Quantification Parameters. Nuclear area was quantified using automated segmentation of DAPI-stained nuclei in Fiji/ImageJ. Individual nuclear measurements were averaged within each field of view and FOV means were subsequently averaged within biological replicates. Statistical comparisons were performed using biological replicate means to avoid pseudoreplication.

| **Parameter** | **Setting** |
| --- | --- |
| Software | Fiji/ImageJ |
| Nuclear marker | DAPI |
| Segmentation method | Automated thresholding and watershed separation |
| Particle size filter | 80-1200 pixels² |
| Circularity filter | 0.30-1.00 |
| Edge objects | Excluded |
| Experimental unit | Biological replicate |
| Intermediate unit | Mean field of view (FOV) |

1	Knerr *et al.* 2001 <https://doi.org/10.1007/s002400000165>

2	Goldberg *et al.* 2007 <https://doi.org/10.1038/sj.jid.5700890>
